# Modeling the benefits of virus discovery and pandemic virus identification

**DOI:** 10.1101/2024.08.26.609801

**Authors:** Geetha Jeyapragasan, Jakob Graabak, Stephen Luby, Kevin M. Esvelt

## Abstract

Preventing and mitigating future zoonotic pandemics are global health priorities, but there are few quantitative estimates of how best to target interventions. Here we construct a mathematical model to evaluate the benefits of 1) virus discovery and sequencing (VDS) in animals and 2) pandemic virus identification (PVI) via laboratory characterization of pandemic potential. Drawing on historical data and expert surveys of One Health and vaccine researchers, we estimate that intensifying virus discovery efforts by three-fold could prevent between 0 and 1.46 million expected deaths per decade by improving non-pharmaceutical interventions and broad-spectrum vaccines. In contrast, because researchers estimate that there are well over a hundred pandemic-capable viruses in nature, identification through laboratory characterization would prevent 48,000 deaths per decade [10,500; 93,600], or just ∼0.62% of expected pandemic deaths. Further identifications would offer diminishing returns. Given wide-ranging survey responses and limited cost-effectiveness compared to proven global health interventions such as insecticide-treated bed nets, our model suggests that health establishments aiming to mitigate future pandemics should focus on monitoring spillover hotspots and empowering local communities to detect, sequence, and suppress nascent epidemics rather than characterizing pandemic potential in laboratories.

## Introduction

Most global pandemics responsible for more than a million deaths have been caused by some form of zoonotic spillover, in which a virus originating in non-human animals jumped into humans, followed by human-to-human transmission, in which the virus or a chimeric descendant spread across the world^1–3^. While many animal viruses might be capable of efficient transmission, only those that spill over into humans can initiate pandemics. In some cases, an initially poorly-transmitting animal virus that infects a human may acquire adaptive mutations or recombine with an endemic human virus to generate descendants capable of efficient transmission, leading to a pandemic^4,5^.

“Disease X” refers to a severe outbreak caused by a currently unidentified pathogen with pandemic potential, which has been placed on lists of priority pathogens for research and development by the World Health Organization and the Coalition for Epidemic Preparedness Innovations^6–8^. As history and epidemiology suggest that zoonotic pandemics are most often caused by RNA viruses ^9,10^, we use the term “Virus X” to represent the hypothetical pathogen responsible for the next pandemic event.

Pandemic prevention efforts aim to reduce the likelihood of Virus X spillover by identifying geographic hotspots and monitoring the animal-human interface, and to interrupt transmission by empowering communities to detect and suppress epidemics before they spread (Fig. 1). Pandemic mitigation efforts aim to develop medical countermeasures and bolster non-pharmaceutical interventions that would be effective against Virus X^6,11^.

**Figure 1.**
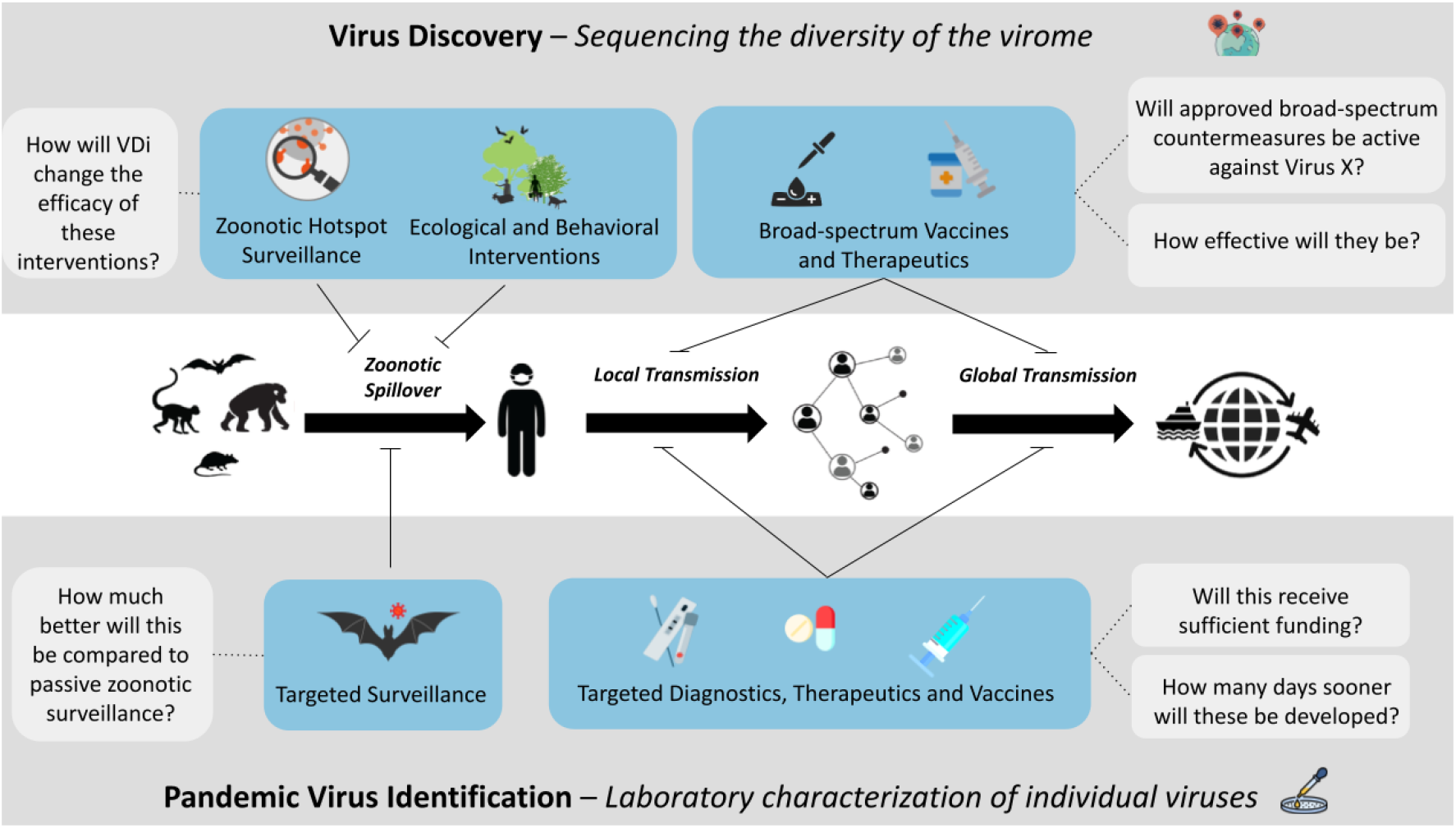
Key steps in a global infectious disease pandemic and candidate interventions. The top panel outlines pathogen-agnostic interventions that could be aided by intensified virus discovery and sequencing (VDS) efforts. The bottom panel depicts targeted interventions that could be enabled or accelerated through pandemic virus identification (PVI) via the experimental laboratory characterization of individual viruses.

Two distinct research strategies attempt to support these prevention and mitigation efforts: Virus Discovery and Sequencing (VDS) and Pandemic Virus Identification (PVI) (Fig. 2).

**Figure 2.**
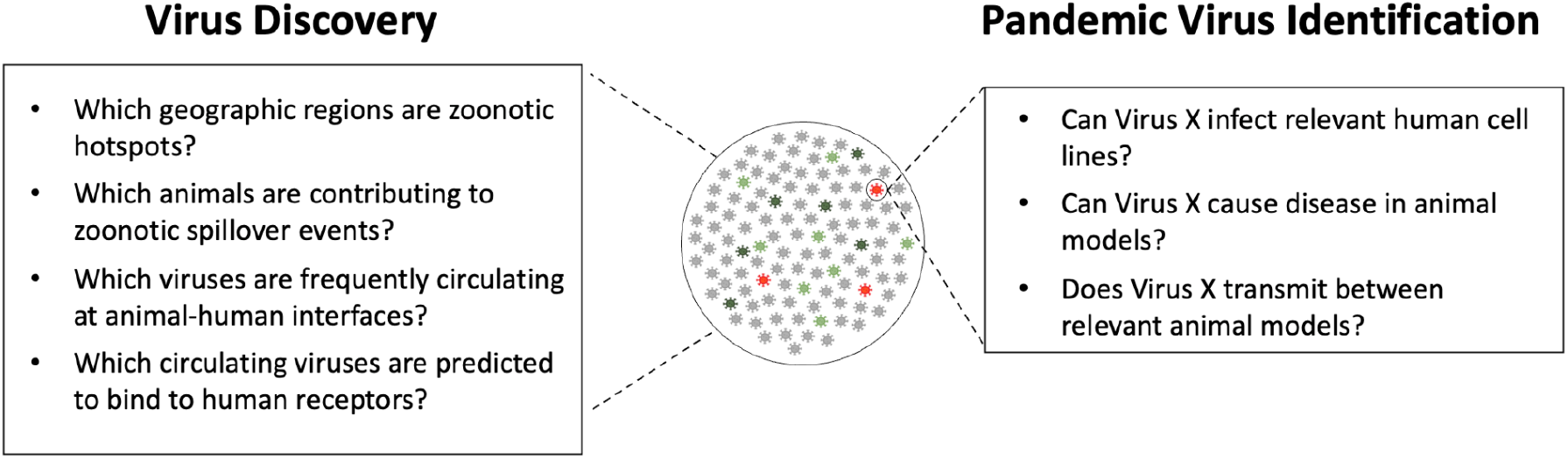
Key questions relevant to virus discovery and pandemic virus identification. Spillover risks can holistically be assessed across pathogens through virus discovery and sequencing efforts (left), while onward transmission risks can be evaluated for individual pathogens through pandemic virus identification efforts (right).

Virus Discovery and Sequencing (VDS) catalogs viral diversity in animal populations to better target anti-spillover and community empowerment interventions and improve the chance that future broad-spectrum vaccines will be effective against Virus X – without needing to identify individual viruses as specific threats^12^. VDS efforts range from those that monitor the animal-human interface in order to catalog those viruses most likely to spill over in humans (community-focused VDS) to sequencing the entire mammalian and avian virospheres, including viruses found in vertebrates remote from human communities (total VDS).

Pandemic Virus Identification (PVI) is a field of research where laboratory experiments are conducted to assess the pandemic potential of individual viruses in human cell cultures and animal models. Initial experiments may include binding and pseudovirus analyses of viral entry proteins, but thorough characterization requires whole-virus infection and replication studies in human primary cells and transmission experiments in animal models^13,14^. While some experiments aim to solely characterize wild-type viruses, others introduce mutations or evaluate potential reassortant viruses to assess whether viruses are within mutational distance of being pandemic-capable^15,16^. Given the complexity of viral infection and transmission in humans, laboratory characterization is required to give any meaningful indication of whether a given virus has the potential to spread efficiently in humans. While developing a targeted vaccine or therapeutic for a virus that has not yet spilled over would be unprecedented, sufficiently concerning results from PVI could in principle unlock sufficient funding^17^ or direct interventions toward communities threatened by identified viruses.

VDS and PVI efforts have often been supported by many of the same researchers and funding programs, such as USAID PREDICT and the Global Virome Project, but they have different goals and methodologies. VDS involves field research in which teams gather bodily fluid samples from animals in zoonotic hotspots and other animal reservoirs, particularly within low to middle income countries. In contrast, PVI experiments are exclusively performed in well-equipped laboratories. Both approaches carry risks: VDS and PVI may risk accidental infections of field researchers or laboratory workers, while PVI may also increase the risk of deliberate misuse by identifying and highlighting novel pandemic-capable viruses.

The financial costs of pandemics are relatively well-established: COVID-19 validated earlier World Bank estimates that moderate to severe pandemics could cause between 14.2 and 71.1 million deaths and global GDP decreases of 2%–4.8%^18,19^. These costs are so large that even slight benefits to prevention and mitigation would be worthwhile. For example, Dobson et al. estimate that as much as $30.7 billion in annual prevention would be cost-effective if investments could accomplish a spillover reduction of 26.7%.

While such funds are unlikely to materialize, they could plausibly support even the costliest VDS and PVI projects. For example, the Global Virome Project aims to sequence 70% of zoonotic viruses for $1.2 billion, or $7 billion for the entire virome (total VDS), then devote additional resources to characterizing the highest-risk viruses and designed mutants that could plausibly exhibit enhanced transmission (PVI)^20^. However, despite extensive research into the factors driving zoonotic risk^7,21–26^, the scarcity of estimates regarding intervention efficacy – let alone empirical data – has precluded quantitative cost-benefit analyses.

There is considerable controversy within virology regarding the potential benefits of VDS and PVI. The debate centers around three main approaches that exist on a continuum:

### Monitoring the animal-human interface (includes some VDS)

Some virologists have asserted that attempting to discover and predict the next pandemic virus is not feasible due to the very large number of viruses in nature ^27,28^. They advocate for monitoring the animal-human interface to better understand spillover risks and spot epidemics earlier, including VDS in at-risk communities.

### Virus discovery and sequencing (VDS)

This strategy extends from community-focused VDS to more comprehensive approaches involving the cataloging and sequencing of viruses in animals far from human populations. Proponents highlight the emerging feasibility of computational methods to spot viruses capable of binding to human receptors, which could increase the likelihood that broad-spectrum medical countermeasures would function against Virus X^29^. Virus discovery in animals living far from humans is somewhat controversial due to the risk of virus hunter infection and onward transmission.

### Pandemic virus identification (PVI)

Advocates of this approach contend that laboratory characterization of viruses capable of human-to-human transmission, or those within mutational reach of this capability (e.g.argue that knowing precisely which viruses can transmit well enough to cause a pandemic, or are within mutational distance of that capability, is crucial for developing vaccines and therapeutics targeting Virus X before it spills over^16^. PVI is controversial due to the risk of laboratory infections and deliberate misuse.

Here we develop a mathematical framework to estimate the anticipated benefits of VDS and PVI. We establish a baseline risk from naturally emerging pandemic events and their sources, estimate the potential reduction in risk from enhanced VDS, then model the further reduction in risk from successful PVI of a single virus and of many viruses. To estimate key parameters for our model and obtain quantitative outcomes, we employed a combination of estimates found in prior literature, close historical proxies to the parameter of interest, and academic surveys sent to domain experts. Given the limited forecasting abilities of experts^30^ and considerable uncertainty in the relevant parameters, we aim to bound the potential benefits from VDS and PVI to inform future public health investments.

## Results

We conducted a survey of experts in One Health and vaccine/therapeutic development to obtain critical parameter estimates for our model. Using historical data and survey responses, we modeled potential benefits of intensifying VDS efforts and successfully identifying pandemic-capable viruses through PVI, using Monte Carlo simulations to account for the wide range of expert opinions and provide 90% central intervals for our median benefit estimates.

### Survey of Experts

Given the dearth of quantitative data concerning key parameters relevant to building a quantitative framework for cost-benefit assessments, we conducted a survey of experts who have repeatedly published in journals from the fields of One Health and/or vaccine and therapeutic development (Methods). Key questions included the likelihood of a broad-spectrum vaccine effective against Virus X with and without virus discovery, the likelihood of funding for and probable acceleration of targeted vaccines and therapeutics given pandemic virus identification, and efficacies and likelihoods associated with non-pharmaceutical interventions (Table 1). In all cases, the gap between the 5th and 95th percentile responses were notably wide, spanning at least 0.55 for every question concerning a probability. We consequently performed Monte Carlo simulations sampling from survey data to translate these uncertainties into 90% confidence windows as potential bounds for the estimated benefits.

**Table 1:**
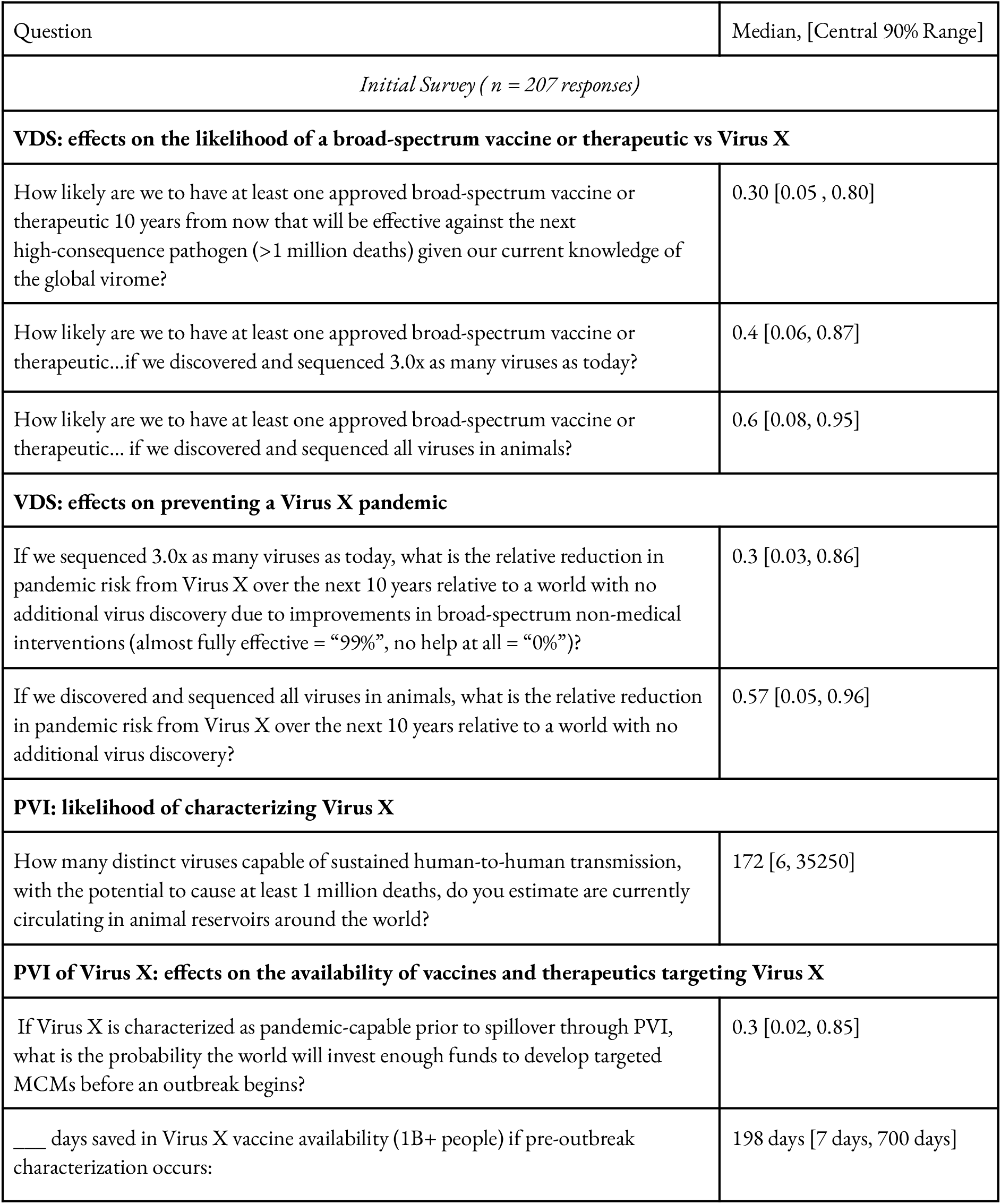
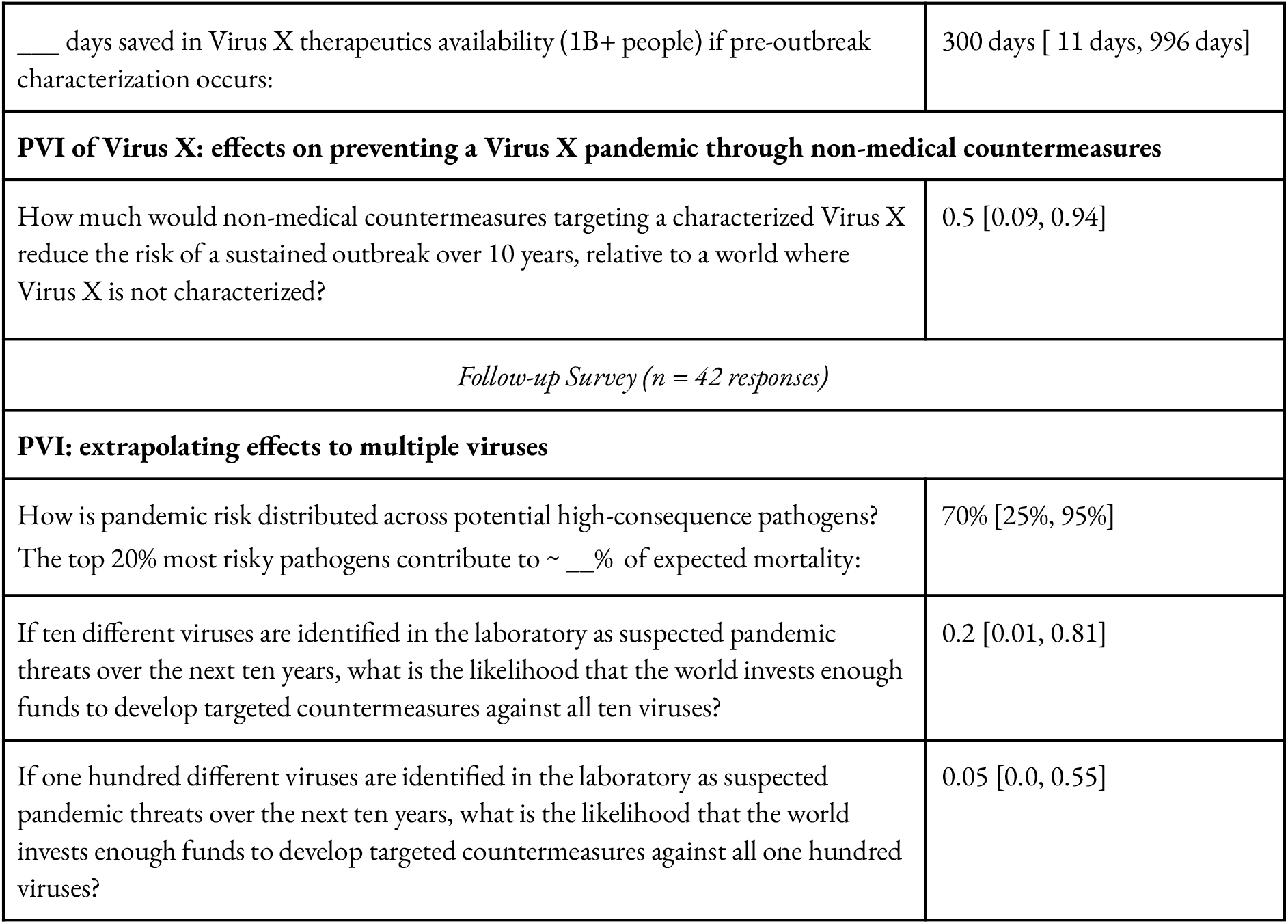
Survey questions and results from the initial survey and follow-up survey.

| Question | Median, [Central 90% Range] |
| --- | --- |
| *Initial Survey ( n = 207 responses)* |  |
| **VDS: effects on the likelihood of a broad-spectrum vaccine or therapeutic vs Virus X** |  |
| How likely are we to have at least one approved broad-spectrum vaccine or therapeutic 10 years from now that will be effective against the next high-consequence pathogen (>1 million deaths) given our current knowledge of the global virome? | 0.30 [0.05 , 0.80] |
| How likely are we to have at least one approved broad-spectrum vaccine or therapeutic...if we discovered and sequenced 3.0x as many viruses as today? | 0.4 [0.06, 0.87] |
| How likely are we to have at least one approved broad-spectrum vaccine or therapeutic... if we discovered and sequenced all viruses in animals? | 0.6 [0.08, 0.95] |
| **VDS: effects on preventing a Virus X pandemic** |  |
| If we sequenced 3.0x as many viruses as today, what is the relative reduction in pandemic risk from Virus X over the next 10 years relative to a world with no additional virus discovery due to improvements in broad-spectrum non-medical interventions (almost fully effective = “99%”, no help at all = “0%”)? | 0.3 [0.03, 0.86] |
| If we discovered and sequenced all viruses in animals, what is the relative reduction in pandemic risk from Virus X over the next 10 years relative to a world with no additional virus discovery? | 0.57 [0.05, 0.96] |
| **PVI: likelihood of characterizing Virus X** |  |
| How many distinct viruses capable of sustained human-to-human transmission, with the potential to cause at least 1 million deaths, do you estimate are currently circulating in animal reservoirs around the world? | 172 [6, 35250] |
| **PVI of Virus X: effects on the availability of vaccines and therapeutics targeting Virus X** |  |
| If Virus X is characterized as pandemic-capable prior to spillover through PVI, what is the probability the world will invest enough funds to develop targeted MCMs before an outbreak begins? | 0.3 [0.02, 0.85] |
| ___ days saved in Virus X vaccine availability (1B+ people) if pre-outbreak characterization occurs: | 198 days [7 days, 700 days] |
| ___ days saved in Virus X therapeutics availability (1B+ people) if pre-outbreak characterization occurs: | 300 days [ 11 days, 996 days] |
| **PVI of Virus X: effects on preventing a Virus X pandemic through non-medical countermeasures** |  |
| How much would non-medical countermeasures targeting a characterized Virus X reduce the risk of a sustained outbreak over 10 years, relative to a world where Virus X is not characterized? | 0.5 [0.09, 0.94] |
| *Follow-up Survey (n = 42 responses)* |  |
| **PVI: extrapolating effects to multiple viruses** |  |
| How is pandemic risk distributed across potential high-consequence pathogens?<br>The top 20% most risky pathogens contribute to ~ ___% of expected mortality: | 70% [25%, 95%] |
| If ten different viruses are identified in the laboratory as suspected pandemic threats over the next ten years, what is the likelihood that the world invests enough funds to develop targeted countermeasures against all ten viruses? | 0.2 [0.01, 0.81] |
| If one hundred different viruses are identified in the laboratory as suspected pandemic threats over the next ten years, what is the likelihood that the world invests enough funds to develop targeted countermeasures against all one hundred viruses? | 0.05 [0.0, 0.55] |

### Virus Discovery and Sequencing

The general lack of international investment in pandemic preparedness even after COVID-19 strongly suggests that virus discovery and sequencing alone is unlikely to ring alarm bells loudly enough to unlock more funding for major interventions ^31–33^. Therefore, researchers expect the main potential benefits to accrue from 1) ensuring that future broad-spectrum medical countermeasures would be effective against Virus X, and 2) improved targeting of existing anti-spillover efforts (Figure 3). Hereafter, we use “VDS” to refer to a scenario featuring a 3-fold increase in the number of viruses discovered and sequenced.

**Figure 3.**
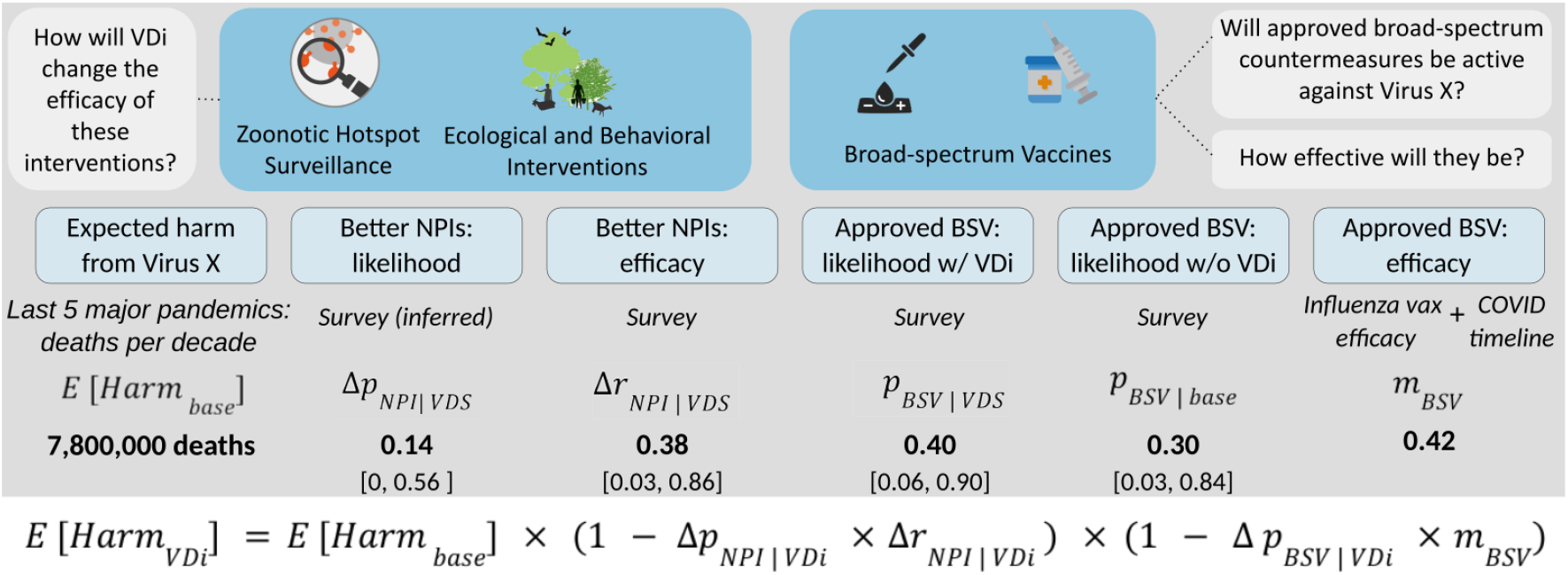
Model and parameters used to estimate harm reduction from virus discovery (VDS). The point value or median used in the baseline model is bolded. Sources of key parameters are indicated. 90% central intervals are available in Tables 1 and 3. Broad-spectrum vaccine efficacy is estimated at half the efficacy of the targeted COVID vaccines, which is approximately that of an influenza vaccine.

Survey participants estimated that the VDS scenario could reduce spillover risk by 30% (Δ*r*_*NPI* | *VDS*_ = 0.3 [0.03, 0.86]) if optimally translated to guide non-pharmaceutical interventions, as current spillover prevention efforts do not take virus density or diversity into account^34^. Achieving the full effect would require that budgets for non-pharmaceutical countermeasures undergo re-allocation based on the findings of the VDS research. Given that updating geographical priorities and re-allocating funds for interventions between nations on the basis of virus discovery and sequencing is presumably more challenging than for the handful of laboratories already developing broad-spectrum vaccines to make use of viral diversity information, we used the estimated relative increase in likelihood of developing an approved broad-spectrum vaccine as an upper bound (Δ*p*_*NPI* |*VDS*_ = 14%; see Methods for higher values).

Respondents assigned a 40% [6%, 90%] probability that a broad spectrum vaccine would be effective against Virus X with VDS (*p*_*BSV* | *VDS*_ = 0.4), as opposed to a 30% [3%, 84%] likelihood of a broad-spectrum vaccine against Virus X without VDS (*p*_*BSV* | *base*_ = 0.3). For the change in likelihood of a broad spectrum vaccine due to VDS, we took the difference (Δ*p*_*BSV* |*VDS*_ = 10% [-13%, 29%]).

To estimate the reduction in harm provided by an immediately available broad-spectrum vaccine against Virus X (*m*_*BSV*_), we calculated the additional deaths that would have been prevented if such a vaccine had already been developed and approved at the start of the COVID-19 pandemic. A 2023 study by Więcek et al. estimated the potential additional lives saved if a targeted COVID-19 vaccine was released earlier in the U.S. and U.K.^35^. Because the Covid-19 vaccines were unusually effective at lowering the risk of death, broad-spectrum vaccines are not expected to be as effective. We consequently assumed that a BSV against Virus X would be as impactful at preventing mortality as the seasonal flu vaccine, which is approximately half as effective as the targeted Covid-19 vaccines. We therefore extrapolated the results of Więcek et al. to estimate the additional reduction in global mortality we would see if a broad-spectrum vaccine for COVID-19 had been available 300 days prior to the approval and release of the targeted vaccines, occurring approximately 50 days into the outbreak. Importantly, we assumed that targeted vaccine development would have proceeded as normal, such that any individuals who received an early broad-spectrum vaccine in our counterfactual would subsequently be vaccinated with a targeted vaccine once it was available. Therefore, most of the benefits from the early availability of the broad-spectrum vaccine would accrue early in the pandemic.

To account for uncertainty in parameter estimates, we conducted Monte Carlo simulations drawing parameter values directly from survey data or distributions derived from the data. These simulations resulted in harm reduction ranging from 0% to 19%, with a median of 9%. This translates to VDS averting 0 to 1.46 million deaths over the following decade, with a median of 492,000 and mean of 741,000. The wide central interval and difference between the mean and median underscores the lack of consensus among experts surrounding potential impacts. Of the expected lives saved, approximately 58% would live because non-pharmaceutical interventions helped to prevent spillover or suppress an outbreak of Virus X, while the remaining 42% would benefit from a broad-spectrum vaccine.

### Pandemic Virus Identification

Given that pandemic virus identification relies on the characterization of individual viruses, we first evaluated the benefits of successful identification of a single pandemic-capable virus.

Survey participants estimated that there are between 5 and 13,395 pandemic-capable viruses in nature, with 70% of the risk concentrated in the top 20% of the median 172 viruses. If we generously assume that the likelihood of a pandemic-capable virus being discovered and subsequently characterized is equivalent to its likelihood of spilling over and causing a pandemic (such that pandemic-capable viruses more likely to spillover are proportionately more likely to be identified), and there is a 3.6% annual probability of a spillover-caused pandemic, then there is a 0.55% chance that any given pandemic-capable virus correctly identified through PVI experiments will seed a pandemic event each decade (Fig. 4).

**Figure 4.**
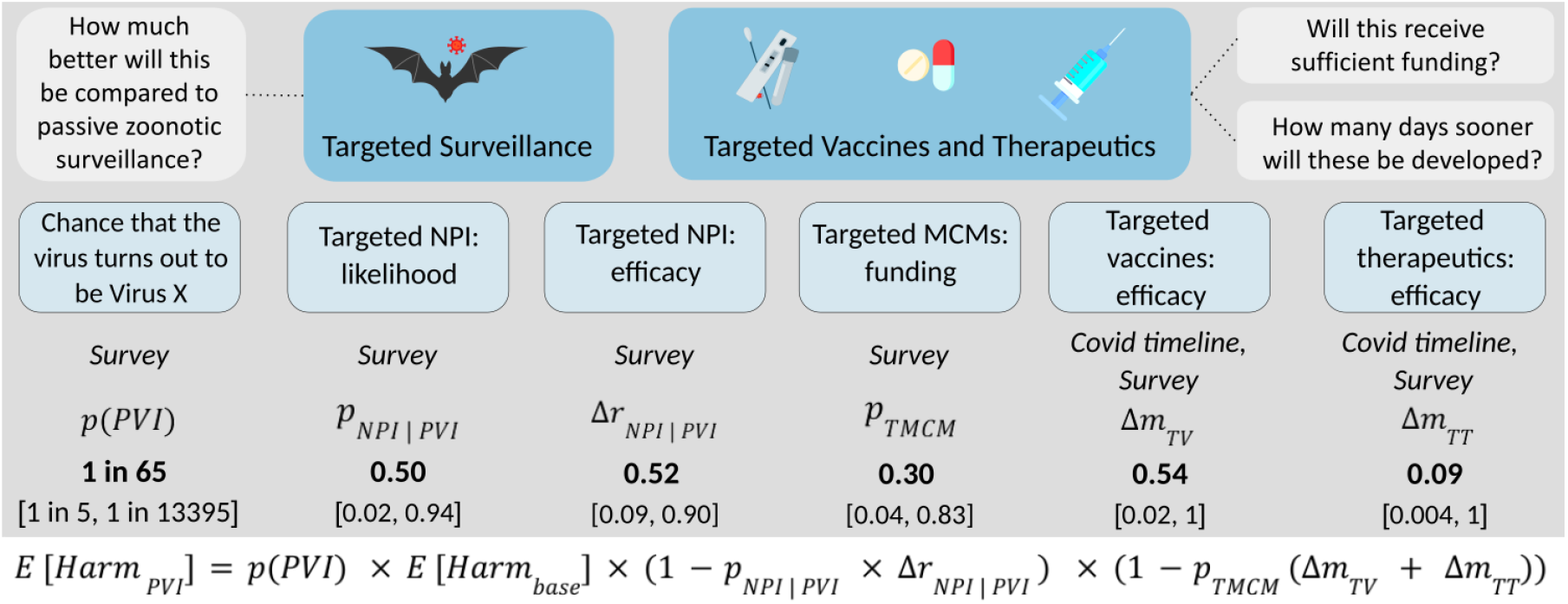
Model and parameters estimating the benefits of pandemic virus identification (PVI). 90% central intervals for each key parameter are provided below each median point estimate. There is a median 0.55% chance that a virus correctly identified through PVI will cause a pandemic in the following decade, which translates to a 1 in 65 chance that the virus would cause the next pandemic if one occurs within that interval. For simplicity, we use this value as a point estimate. For Δ*m*_*TV*_, participants estimated a vaccine would be accelerated by 7 to 700 days, and for Δ*m*_*TT*_, that a therapeutic would be accelerated by between 10 and 995 days. Since we used COVID-19 timelines and modeling by Więcek et al. as a base to convert these accelerations into relative increases in lives saved, and SARS-CoV-2 vaccines and therapeutics were available 357 and 772 days after pandemic onset, the upper bounds of the central intervals would be greater than 1, paradoxically saving more lives than would have perished in the pandemic. We therefore cut off the upper bounds at 1.

We grouped the benefits of PVI into two categories: the benefits of targeting NPIs toward regions at high risk of identified virus spillover, and the benefits it would provide to the production of medical countermeasures through accelerating timelines.

We first sought to estimate the reduction in pandemic risk achieved by characterizing Virus X and targeting interventions to prevent spillover and control outbreaks. Participants estimated that if the virus in question was in fact Virus X, better-targeted interventions would reduce spillover risk by an additional 60% [9% to 94%]. We multiplied these benefits by likelihood that a discovered virus subjected to PVI would cause a pandemic within the subsequent decade (0.55%).

To assess the extent to which medical countermeasures would be accelerated, participants were asked to estimate how much earlier a vaccine and therapeutic for Virus X would be available to at least 1 billion people if the virus had been characterized and flagged as a suspected pandemic risk before the outbreak. The median response indicated that *if* sufficient funding were acquired, a targeted vaccine would be available 198 days sooner [7 days, 700 days], and a therapeutic would be available 300 days sooner [11 days, 996 days], compared to a scenario where Virus X had not been preemptively identified. Combining these results with COVID-19 medical countermeasure data, we estimated these accelerated timelines would save 50% more lives through earlier vaccination and 9% more from earlier therapeutic availability relative to a world in which Virus X had not been identified in advance, assuming funding. When asked to estimate the probability that targeted vaccines and therapeutics would receive sufficient funding if Virus X were to be characterized, the median participant response was 30% [2%, 85%]. To calculate the net benefits, we multiplied the estimated lives saved by the estimated probability of funding and the likelihood that the virus characterized would be Virus X.

Given the wide range of survey responses, we conducted Monte Carlo simulations sampling from survey data to represent the parameters *p*_*TMCM*_, *p*_*NPI* | *PVI*_, and Δ*r*_*NPI* | *PVI*_. These simulations estimated that successful PVI would save between 10,500 and 93,600 lives (90% central interval), with a median benefit of approximately 48,000 lives saved.

Next, we considered how the benefits scale with each additional successfully identified virus. In our follow-up survey, experts estimated the likelihood of sufficient funding being pooled for targeted countermeasures, including NPIs, vaccines, and therapeutics, if multiple pandemic viruses were identified. Monte Carlo simulations revealed that identifying all 172 pandemic-capable pathogens in nature would reduce natural pandemic risk by between 0.026% and 28.7% over the next decade, saving between 4,600 and 5.1 million lives in expectation (median 48,000; Figure 5).

**Figure 5.**
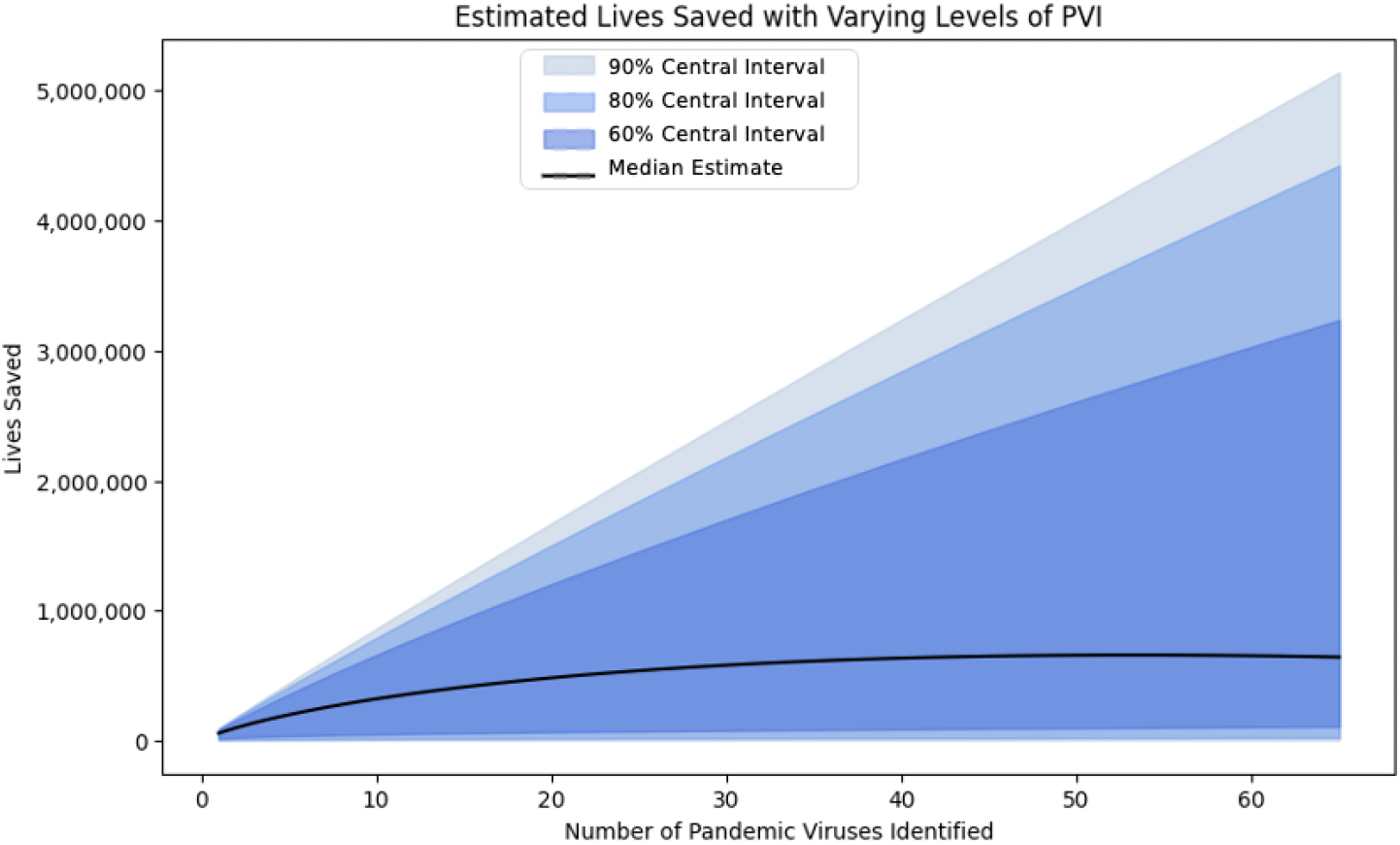
Estimated number of lives saved as a function of pandemic-capable viruses identified. The black line represents the model outputs using the median *p*_*TMCM*_ (and *p*_*NPI* | *PVI*_) estimate, and the bands represent model outputs using the 60%, 80% and 90% central intervals of the parameter.

Notably, identifying all pandemic-capable viruses would require researchers to discover and sequence all viruses and conduct laboratory characterization experiments on each computationally predicted high-risk pathogen to assess its pandemic potential.

### Comparison of VDS and PVI

A three-fold increase in VDS is expected to save zero to 1.46 million lives [median 492,000] over the next decade. PVI of a single virus is expected to save between 10,500 and 93,600 [median 48,000], or 4,600 to 5.1 million if every pandemic-capable virus in nature [median 172] was successfully identified through characterization (Fig. 6). Crucially, the latter level of PVI would require a greater investment in VDS than the threefold increase that we evaluate, as only a discovered virus can be characterized. These results are heavily dependent upon parameters from the expert surveys, which exhibited considerable variance.

**Figure 6.**
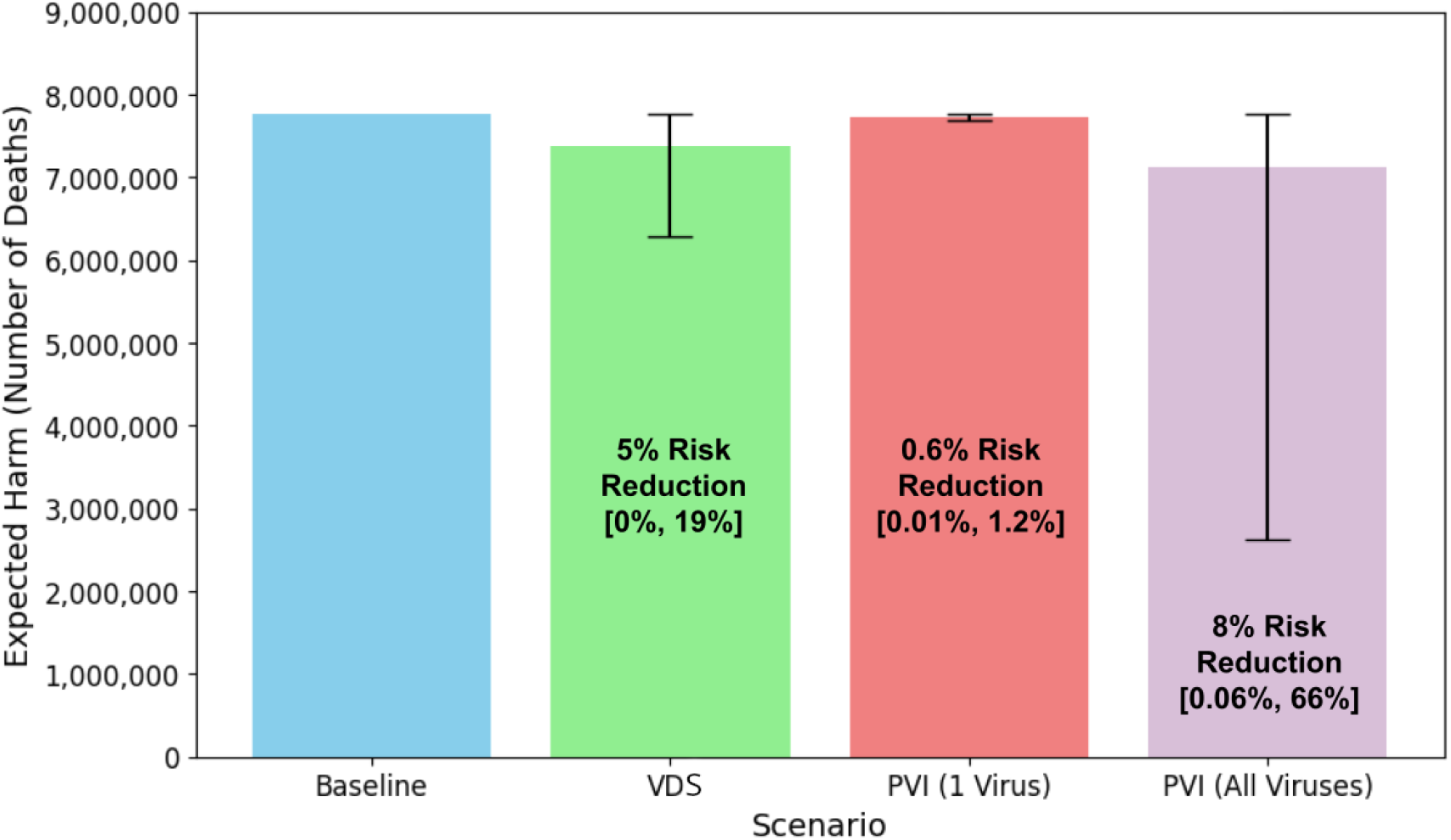
Comparison of expected harm from a Virus X pandemic over the next decade. The baseline scenario depicts the status quo, the VDS scenario features the discovery and sequencing of 3 times as many viruses as today, the PVI (1) scenario depicts the results of characterizing the first presumably pandemic-capable virus, and the PVI (all) scenario characterizes all pandemic-capable viruses in nature. The error bars represent the 90% central interval for each estimate.

## Discussion

Our benefit estimation framework and expert survey provide insights into the potential benefits of viral discovery efforts and pandemic virus identification (PVI) research in mitigating future pandemic risks.

The model suggests that discovering and sequencing three times as many viruses could save between 0 and 1.46 million lives from a Virus X pandemic in the next decade, with a median of 492,000. These benefits arise from the hypothetical ability to better target NPIs to regions with high viral diversity and the increased likelihood that broad spectrum pharmaceuticals would be effective against an emerging Virus X pandemic.

In contrast, pandemic virus identification would save an estimated 10,500 to 93,600 [median 48,000] lives per virus identified in the next decade, scaling to 4,600 to 5.1M [median 642,000] lives saved per decade given successful discovery and characterization of all ∼172 [6, 35250] pandemic-capable viruses in nature. Most of these benefits would accrue from accelerating targeted medical countermeasures.

A key limitation of this study is its dependence upon parameters obtained from surveys of domain experts, who returned an extremely broad range of estimates for virtually all probabilistic questions, and may suffer from systematic biases. The work of Tetlock and coworkers^30^ has demonstrated that in the field of political judgment and foreign relations, even highly qualified individuals struggle to make accurate long-term predictions, especially on complex topics. This variance is reflected in high uncertainties.

Despite contested claims over the benefits of virus discovery and characterization efforts, participant views were largely unaffected by field^27,36,37^. Experts working in virus discovery and characterization programs estimated a median 10% increase in the probability of a broad-spectrum vaccine given intensified VDS, versus 7% to 9% in other groups (national academies, basic research and applied research cohorts), and were also somewhat more optimistic about the probability of targeted medical countermeasures receiving sufficient funding (40%) given PVI relative to the other groups (20 to 30%). Nevertheless, estimates for vaccine and therapeutic timeline acceleration were quite varied across groups, with Basic Research providing the highest estimates for vaccine acceleration (200 days). Relative to the wide ranges reported overall, these field-specific differences are minor.

High expert uncertainty and potential systemic biases may instead reflect the speculative nature of many potential outcomes, some of which are heavily influenced by factors beyond their areas of specialty. For example, funding support to develop a vaccine for a virus that has not yet infected a human would be unprecedented and therefore surprising. Nipah virus – a highly lethal potential pandemic pathogen which has caused multiple outbreaks exhibiting sustained human-to-human transmission since 1998 ^38^ – still has no approved medical countermeasures, and only recently have candidate vaccines entered clinical trials^39^. Yet experts assigned a 30% [4%, 83%] chance of this occurring, perhaps viewing the recent funding for Nipah vaccines as evidence of progress.

That PVI-triggered funding might accelerate targeted vaccine timelines by hundreds of days would be similarly surprising given that the first mRNA vaccine entered Phase 2 trials just 140 days after publication of the genome sequence – and due to the prototype pathogen approach^40^, in which a vaccine is developed against one member of each viral family, future targeted vaccines against Virus X are expected to enter combined Phase 1/2 trials immediately. Pre-existing manufacturing plants are expected to enable still more rapid production^40^. Survey participants may have assigned a high probability of mRNA vaccines and similarly rapid adenoviral vaccines being ineffective against Virus X, or anticipated a slower-spreading outbreak than those of the influenza and coronavirus pandemics of the past 135 years.

Therefore, while our mathematical model provides a structured approach to estimating potential benefits, the results of this study should be viewed as an exploratory attempt to bound potential outcomes, not as definitive benefit assessments. Future research should focus on refining key model parameters as more empirical data becomes available. In the interim, our results may provide guidance for funders considering how best to allocate scarce pandemic prevention and mitigation resources in a cost-effective manner.

For example, judging by the fact that sequencing 70% of mammalian and avian viruses was estimated to cost $1.2 billion in 2018^41^, the VDS scenario may cost several hundred million to a billion dollars today. Our model, informed by the expert surveys, predicts that such an investment could save an estimated median of 492,000 [0, 1.46M] lives from a Virus X pandemic in the next decade, with the caveats of high uncertainty and the unprecedented nature of several key answers. Using the median estimate as a point of comparison such an investment would be only somewhat more cost-competitive than increased spending on well-evidenced areas of public health such as anti-malaria bed nets ^42–44^. However, given the large amounts of uncertainty associated with the benefits of VDS, allocating limited funds towards interventions with less uncertainty may be preferable.

The cost to identify a single pandemic-capable virus is unknown because the accuracy of computational prediction is not known, but identifying approximately all pandemic-capable viruses in order to save ∼642,000 lives over a decade would first require discovering and sequencing the entire mammalian and avian viromes, estimated at $7 billion^41^, then performing expensive characterization experiments involving human primary cell infection and replication and animal transmission on hundreds or thousands of candidate viruses. However, it may not be reasonable to compare anti-pandemic interventions with other forms of global health interventions. Investments into superior protective equipment or germicidal lights may be more appropriate comparators, although such research has not yet been subjected to quantitative benefit estimates or even expert surveys. Unlike PVI, pathogen-agnostic investments do not carry dual-use risks.

Perhaps the most appropriate alternative to investing more funds into VDS and PVI is one that was also funded by USAID PREDICT: the empowering of communities in zoonotic hotspots to prevent spillover and suppress epidemics before they can spread. Thanks to recent advances in biotechnology, these efforts could be much more effective than when PREDICT began. Local communities with access to nanopore sequencing technology can obtain and share the sequence of a novel pathogen within a day of recognizing an outbreak^45^, potentially enabling the development, manufacturing, and delivery of CRISPR-based rapid diagnostics and targeted nucleic acid vaccines within weeks or even days^46^. Deploying these in a combined Phase 1/2 vaccination trial, including ring vaccination surrounding anyone ill who tests positive, could maximize the likelihood of containing the outbreak^47^. In a “1-10-10k” plan, a genome sequence would be made available to the world on day 1 of the epidemic being recognized, and ten days later, the world would deliver 10,000 rapid diagnostic tests and 10,000 doses of nucleic acid vaccine for targeted ring vaccination. As with VDS and PVI, such a proposal could benefit from a quantitative cost-benefit assessment.

Collectively, our results suggest that despite the high uncertainty in key parameters obtained from expert surveys, quantitative modeling of the expected benefits and risks of proposed public health research programs can translate the collective views of the scientific community into outer-bound estimates relevant to deciding how best to allocate scarce resources.

## Methods

To construct the model, we first surveyed the literature to gather a list of pandemic prevention interventions, which we group into three broad categories: preventing initial spillover from animals to humans, suppressing transmission to prevent end nascent epidemics, and mitigating harms from an epidemic that has spread to become a global pandemic (Table 2) ^48^. Through this review, we noted which interventions researchers listed as those VDS and PVI could potentially influence through providing information that could influence prioritization or directly contribute information valuable for countermeasure development.

**Table 2:** Pandemic mitigation interventions.

|  | Category | Intervention | VDS | PVI |
| --- | --- | --- | --- | --- |
| **Preventing spillover**<br>(Primary prevention) | Research | Map zoonotic hotspots | ✓ |  |
|  |  | Monitor high-risk animal reservoirs | ✓ | For Virus X |
|  | Land-use | Prevent deforestation near hotspots | ✓ |  |
|  |  | Minimize habitat fragmentation | ✓ |  |
|  | Markets | Strictly regulate and monitor the wildlife trade | ✓ | For Virus X |
|  |  | Strictly regulate and monitor wet markets | ✓ | For Virus X |
|  |  | Improve animal husbandry practices | ✓ | For Virus X |
|  | Education | Educate communities about risks from animal reservoirs | ✓ | For Virus X |
|  |  | Offer training in safe practices for at-risk occupations | ✓ | For Virus X |
| **Suppressing transmission**<br>(Secondary prevention) | Diagnostics | Develop targeted rapid tests for specific viruses |  | For Virus X |
|  |  | Develop rapid tests for high-risk virus families | ✓ |  |
|  |  | Equip communities with sequencing-based diagnostics | ✓ |  |
|  |  | Develop and administer broad-spectrum vaccines | ✓ |  |
|  | Health workers | Train healthcare providers to use diagnostics and vaccines | ✓ |  |
|  |  | Provide protective equipment and safety training | ✓ |  |
|  | Containment | Preparing for travel restrictions in the event of an epidemic | ✓ |  |
| **Mitigating harm** |  | Develop and approve targeted vaccines |  | For Virus X |
|  |  | Develop and approve broad-spectrum vaccines | ✓ |  |
|  |  | Develop and approve broad-spectrum therapeutics | ✓ |  |
|  |  | Develop and approve targeted therapeutics |  | For Virus X |

*\* Relevance of PVI determined by assessing current measures introduced to prevent a Nipah virus pandemic*

All interventions would benefit from improved resource allocation towards higher-risk communities, reservoirs, and pathogens. VDS can increase the likelihood that any broad-spectrum vaccines will be effective against Virus X, and may improve hotspot targeting, which currently relies on biodiversity estimates. PVI allows for the development of targeted pathogen-specific vaccines and therapeutics and may direct anti-spillover efforts towards regions with identified pathogens. Theoretically, the availability of vaccines or therapeutics for a virus that has successfully begun transmission between humans could prevent an uncontrolled outbreak through ring vaccination efforts, both reducing the chances of a local outbreak spreading globally, and mitigating the amount of morbidity and mortality caused by outbreaks.

We first established a baseline risk of pandemics over the next decade, evaluating the expected harm of natural zoonotic pandemics based on the likelihood of occurrence and the consequences of a pandemic (measured by the average number of deaths posed by the pathogen over the next decade). Then using the estimated number of pandemic viruses circulating around the world by survey participants, we estimated the expected harm posed by the next virus to cause a pandemic - Virus X. We use risk synonymously with expected harm, where:

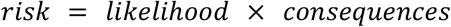

To evaluate the benefits of VDS, we considered the scenario where VDS efforts discover and sequence 3 times as many viruses as we know today through metagenomic sequencing efforts.

We then consider the scenario where Virus X is discovered and identified as a pandemic-capable virus. In this scenario, laboratory experiments are conducted to assess whether primary human cell lines are permissive to efficient infection and amplification of Virus X, and whether Virus X is transmissible in relevant animal models (e.g. ferrets, humanized mice). The results of these experiments, alongside the whole genome sequence of Virus X, are made publicly available to be used in surveillance and potentially used for MCM development efforts.

To obtain key parameters for the model, we used a combination of estimates found in prior literature, close proxies to the parameter of interest, and an academic survey sent to domain experts. The inclusion criteria to select domain experts is outlined in the parameter estimation section below.

### Introduction to Scenarios

This model establishes a framework to estimate the per decade expected pandemic harm from a *Virus X* pandemic in three scenarios: the “baseline” scenario, the virus discovery scenario, and the pandemic virus identification scenario. There are numerous ways in which the world could look with and without VDS or PVI. The assumptions we made about what each of these hypothetical scenarios look like with respect to research, prevention and response activities of Virus X.

#### Baseline Scenario

- Virus X remains unknown to any humans and only exists in the zoonotic hotspot prior to spillover.
- Virus X’s presence is only revealed if X spills over into the human population. The initial of humans infected with Virus X would result in a cluster of atypical symptoms, alerting health agencies to the novel threat, resulting in X being identified and sequenced. Interventions to prevent zoonotic spillover of Virus X are typical threat-agnostic interventions carried out in all high-risk interfaces without knowledge of a specific threat.
- Development of any Virus X-specific therapeutic or vaccine begins after detection in humans. We assume the development timelines will mirror those of the recent COVID-19 pandemic. Additionally, we anticipate it would take 722 days to develop a targeted antiviral, based on the timeline for Paxlovid’s development and approval.

#### Virus discovery (VDS) Scenario

- Samples from animals within a high-risk zoonotic hotspot are collected. These samples are transported back to a lab and sequenced, potentially resulting in the discovery of Virus X (amongst several other pathogens).
- Virus X is potentially amongst the several virus species discovered, but is not prioritized or flagged as a pathogen of special concern beyond computational predictions. If Virus X is in a viral genus/family that contains other viruses that have demonstrated epidemic/pandemic potential (e.g. influenza, coronaviruses, filoviruses, paramyxoviruses), broad spectrum therapeutics and vaccines might be developed for the viral family/genus of Virus X that otherwise would not have covered Virus X.
- If Virus X is circulating in a hotspot deemed to be particularly high priority, either due to its genus or other viruses found in the region, the hotspot may be prioritized for non-pharmaceutical interventions (see Table 2 for examples).

#### Pandemic Virus Identification (PVI) Scenario

- A series of characterization experiments are conducted to estimate the probability Virus X has pandemic potential. The whole genome sequence of Virus X and the results of characterization experiments are publicly published and distributed to relevant stakeholders.
- This information <u>potentially</u> results in targeted non-medical interventions (e.g. Virus X specific surveillance where Virus X was found), as well as efforts to develop promising therapeutic and vaccine candidates for Virus X prior to spillover.

## Mathematical Model

### Parameter Estimation

Quantifying the public health benefits of both viral discovery and PVI research efforts can be quite challenging, both due to the inherent challenges associated with evaluating the benefits of any form of scientific research, and the lack of research evaluating the efficacy of various pandemic prevention strategies.

In constructing our mathematical model to assess the benefits of both VDS and PVI, we first used prior literature on historical pandemics as well as data from the recent COVID-19 pandemic to establish a baseline risk from natural pandemics, as well as establish baselines for targeted vaccine and therapeutics efficacies, as well as development and distribution timelines, distribution timelines and efficacies.

We also identified several key parameters that had not been estimated in prior literature. To gather estimates for these parameters, we sent out two surveys to experts in relevant fields. Participants for the survey were selected based on a criteria established around academic journal publications. Specifically, we identified all authors who had published at least twice between Jan 1, 2019 and May 1, 2023 in any of the following journals: *The Lancet Infectious Diseases, Emerging Infectious Diseases, Immunity*, and *Nature Reviews Immunology*. The publications had to be either in the ‘Article’ or ‘Review’ category, and could not have more than 30 authors. The two publications did not necessarily need to be in the same journal. This provided us with a list of 3,557 authors. We sent out two surveys, an initial survey asking participants to estimate various parameters of our model (n=207), and a followup survey to gather clarification for specific parameters where there was ambiguity, and estimate additional parameters (n = 42). Table 3 outlines some key parameters of the mathematical model, noting which parameters we were able to generate estimates for using prior literature, and which required input from relevant experts.

**Table 3:**
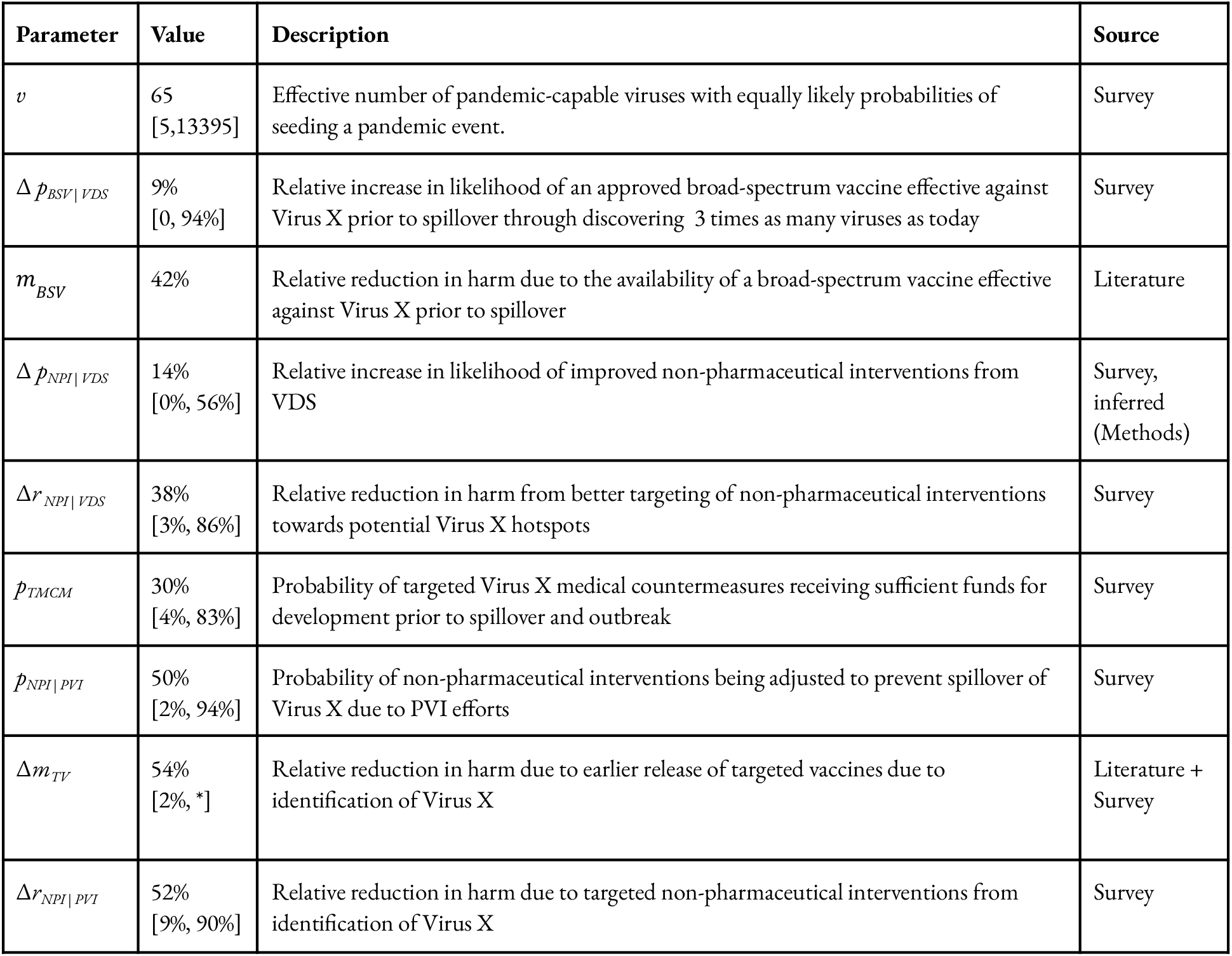
Key Model Parameters. *Complete table of parameters found in supplementary document

The complete parameter table, survey data and data used to estimate the remaining parameters can be found in the supplementary document. The * was used to denote where the upper bound estimate was greater than 100% due to participants’ upper bound estimates of accelerated vaccine and therapeutics development timelines, which would have resulted in more lives saved than those lost in pandemic events.

#### Baseline Risk

Since 1889, five natural pandemics have killed over a million people within a few years of spilling over. Historical data suggests a 3.75% annual likelihood of a natural pandemic event, with an average severity of 18.1. million deaths. These results closely match those of Fan et al., who estimate the overall annual probability of a pandemic to be 3.6% with an average severity of 21.6 million deaths ^49^. For simplicity, we use their figures for subsequent calculations.^15^ This results in approximately 7.8 million expected deaths per decade from zoonotic pandemics, underscoring the importance of effective interventions.

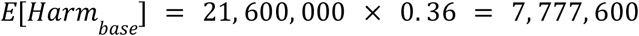

To establish the baseline harm from a Virus X pandemic, we make the simplifying assumption that the pandemic risk over the next decade is from a novel virus rather than an already-known pathogen such as Nipah. Accordingly, the baseline harm from a Virus X pandemic over the next decade is 7.8 million.

#### Virus Discovery

We define VDS as the scenario where three times as many viruses are discovered in high-risk hotspots as we know today. Through our model, we aim to answer the question: “How would the risk of a Virus X natural pandemic event decrease if we discovered and sequenced three times as many viruses as we have today through current discovery and monitoring efforts?”

Our model considers two pathways in which VDS could cause downstream changes to reduce the risk of a Virus X pandemic. The first path is through the influence of pathogen-agnostic or broad-spectrum non-pharmaceutical interventions (NPIs), where the Virus X hotspot(s) might be prioritized for some spillover prevention interventions (Table 2). The second pathway is through influencing the development of a pan-genus or pan-family broad-spectrum vaccine (BSV) that otherwise would either not have been developed and approved, or would not have worked against Virus X. For each pathway, we consider both the probability that VDS will influence the efficacy of the intervention, and the change in efficacy of the interventions themselves.

We consider a few key parameters to quantify the reduction in risk from VDS. First, we consider the change in likelihood that BSV will be effective against Virus X due to VDS (Δ*p*_*BSV* | *VDS*_). We also consider the magnitude of the reduction in harm a broad-spectrum vaccine would provide if available at the start of the outbreak (*m*_*BSV*_).

For non-pharmaceuticals, we note the key parameters as the difference in likelihood of prioritized non-pharmaceutical interventions due to VDS efforts (Δ*p*_*NPI* | *VDS*_) and the reduction in harm from prioritized non-pharmaceutical interventions due to VDS (Δ*r*_*NPI* | *VDS*_). Using these parameters, we define the reduction in pandemic risk from Virus X due to VDS as follows, and walk through the calculations using the median estimate of each parameter:

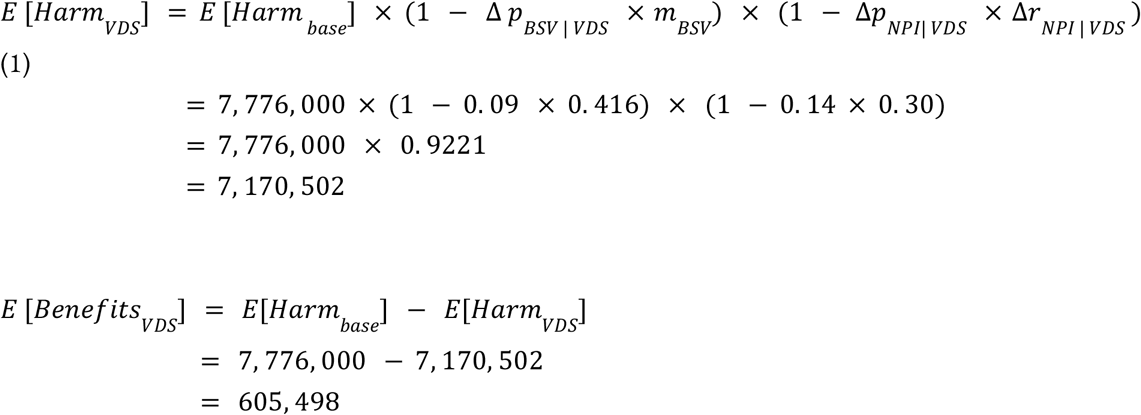

Due to uncertainty over the effectiveness of NPIs and levels of future investment, we make the simplifying assumption that the probability of NPI efficacy against Virus X given VDS (Δ*p*_*NPI* | *VDS*_) is equal to the required relative increased probability that a BSV is developed for Virus X due to VDS (Δ*p*_*BSV* | *VDS*_). Increasing the probability of a broad-spectrum vaccine from a median of 0.3 to a median of 0.4 requires a 14% increase in the overall likelihood of success: (0.40 – 0.30) / (1 – 0.30). We use survey results to estimate Δ*p*_*BSV* | *VDS*_ and Δ*r*_*NPI* | *VDS*_ and estimate the value of *m*_*BSV*_, extrapolating data based on a 2023 study by Więcek et al. evaluating the potential benefits if COVID-19 vaccines had been available earlier in the outbreak (see supplementary document for derivation).

The calculation above uses the median of each parameter as a point estimate. Due to the majority of the parameters being risk-skewed, the mean point estimate is greater than the median generated by our Monte Carlo simulations which directly sample from the survey data, where the mean benefit is 740,000 lives saved and the median benefit is approximately 492,000 lives saved. To reduce the skew from a handful of very high responses, we report the median value from the Monte Carlo.

## Uncertainty Quantification

The calculations above use the mean as point estimates for parameters based on survey data, though there is a large amount of uncertainty amongst experts reflected in the wide distributions of the various questions. To account for uncertainties in our parameter estimates, we conducted Monte Carlo simulations using Python with the NumPy and SciPy libraries. We performed 100,000 iterations for each analysis, drawing parameter values directly from survey data and from existing literature review. For each iteration, we calculated the harm reduction and deaths averted using our model equations, generating distributions of possible outcomes. From these distributions, we computed 90% central intervals as well as means and medians to characterize the bounds and uncertainties in our results.

### Scenario Analysis

In the initial model, we make the assumption that the change in likelihood of NPIs being targeted towards preventing Virus X, Δ*p*_*NPI* | *VDS*_ ; is equivalent to the change in likelihood of a broad-spectrum vaccine being approved for use against virus X, Δ*p*_*BSV*| *VDS*_. Here, we relax that assumption and consider scenarios where they are not equivalent, and run MC simulations setting Δ*p*_*NPI* | *VDS*_ to 30% and 50%, running 100,000 simulations at each level. We report these results in table 4 below.

**Table 4:** Results of MC Simulations at Various Levels of Δ*p*_*NPI* |*VDS*_.

| $\Delta p_{NPI\vert VDS}$ | Lives Saved |
| --- | --- |
| 30% | $\bar{x}$: 1,129,123<br>M: 1,075,452<br>90% central interval: (69,884 ; 2,235,009) |
| 50% | $\bar{x}$: 1,695,476<br>M: 1,555,200<br>90% central interval: (161,741 ; 3,341,752) |

#### Pandemic Virus Identification (PVI)

To evaluate the benefits of pandemic virus identification, we aim to evaluate the question “How would the risk of a natural Virus X pandemic decrease if we identified Virus X as a pandemic capable virus prior to spillover?”. For the purposes of this model, we make a few simplifying assumptions:

1. There are a set number of *pandemic capable* viruses circulating around the world
2. Pandemic virus identification efforts conducted for a given pathogen will reveal with whether the virus is pandemic capable through characterization experiments estimating the virus’s virulence and transmissibility in humans

The primary proposed benefits of PVI that are not possible with VDS alone include targeted pathogen specific medical countermeasures, and targeted non-pharmaceutical interventions. Knowledge that Virus X is a pandemic-capable virus may result in prioritization of Virus X spillover prevention and mitigation efforts, leading to greater allocation of resources towards this specific threat, and an earlier start to development of vaccines and therapeutics. Based on the key potential benefits noted in literature about PVI, we chose the following parameters to quantify the additional reduction in risk: the likelihood a targeted MCM is funded following identification (*p* _*TMCM*_), reduction in harm through earlier release of targeted vaccines (Δ*m* _*TV*_) and therapeutics (Δ*m* _*TT*_) due to PVI, the likelihood PVI results in changes to threat-agnostic interventions and the relative reduction in pandemic risk from PVI informed non-pharmaceutical interventions (Δ*r* _*PVI, NPI*_).

Given that pandemic virus identification relies on the characterization of individual pandemic viruses, we first evaluate the pandemic risk posed by an individual pandemic virus. Through our survey, we estimate a 0.55% chance that any given identified pandemic-capable virus will prove to be Virus X. This is because there are the equivalent of 65 [5; 13,395] pandemic-capable viruses that are equally likely to seed a pandemic event (see Supplementary Information for derivation of the likelihood and equivalent number of pandemic viruses estimates). It should be noted that to date, no viruses identified as potential pandemic pathogens without spilling over into humans have resulted in the development of targeted interventions. This model starts with the assumption that PVI has successfully characterized a novel zoonotic pandemic-capable virus before spillover, and first estimates the benefits of PVI per successfully identified pandemic virus. We first establish the probability PVI has successfully characterized Virus X, rather than a different pandemic-capable virus, as:

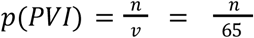

where *n* is the number of viruses characterized and *v* is the number of pandemic-capable viruses that are equally likely to seed a pandemic event.

Based on the key potential benefits noted in literature about PVI, we chose the following parameters to quantify the additional reduction in risk: the likelihood a targeted MCM is funded following identification (*p*_*TMCM*_), the reduction in harm through earlier release of targeted vaccines (Δ*m*_*TV*_) and therapeutics (Δ*m*_*TT*_) due to PVI, the likelihood PVI results in changes to threat-agnostic interventions (*p*_*NPI* | *PVI*_), and the relative reduction in pandemic risk from PVI informed non-pharmaceutical interventions (Δ*r*_*NPI* | *PVI*_).

We define the expected harm in a scenario with PVI as follows:

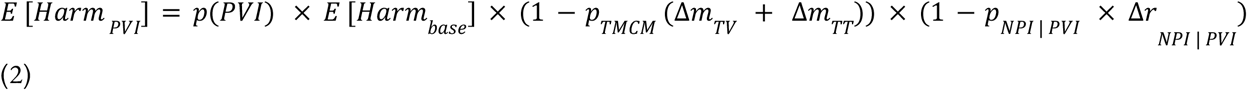

For the parameters above, *p*_*MCM*_ was estimated through the survey, where participants were asked to estimate the likelihood discovering Virus X would lead to sufficient funding being pooled to develop targeted medical countermeasures. To generate estimates for Δ*m*_*TV*_ and Δ*m*_*TT*_, we estimated both how much earlier targeted vaccines and antivirals would be released, as well as the efficacy of the MCM itself. To estimate the shortened timeline, we asked survey participants how much earlier they anticipate a targeted vaccine and a targeted therapeutic would be released if PVI identified Virus X as a pandemic-capable virus, providing an estimate of the number of days. We also used the efficacies and distribution timelines of COVID-19 vaccines and antivirals following emergency use authorization and additional modeling literature to estimate the potential additional lives saved due to earlier release of these interventions in this scenario. For the case of a single pandemic virus successfully identified, we estimate this could result in an average approximately 48,000 lives saved in expectation.

### Scaling Up PVI

To estimate the benefits of the entire PVI enterprise, we considered how the benefits scale for each additional identified virus. In our followup survey, we asked participants about the likelihood of sufficient funding being pooled if multiple pandemic viruses were identified. First using the median estimates of participant answers, we plotted the likelihood of acquiring sufficient funds against the number of viruses successfully identified (Fig. 7).

**Figure 7.**
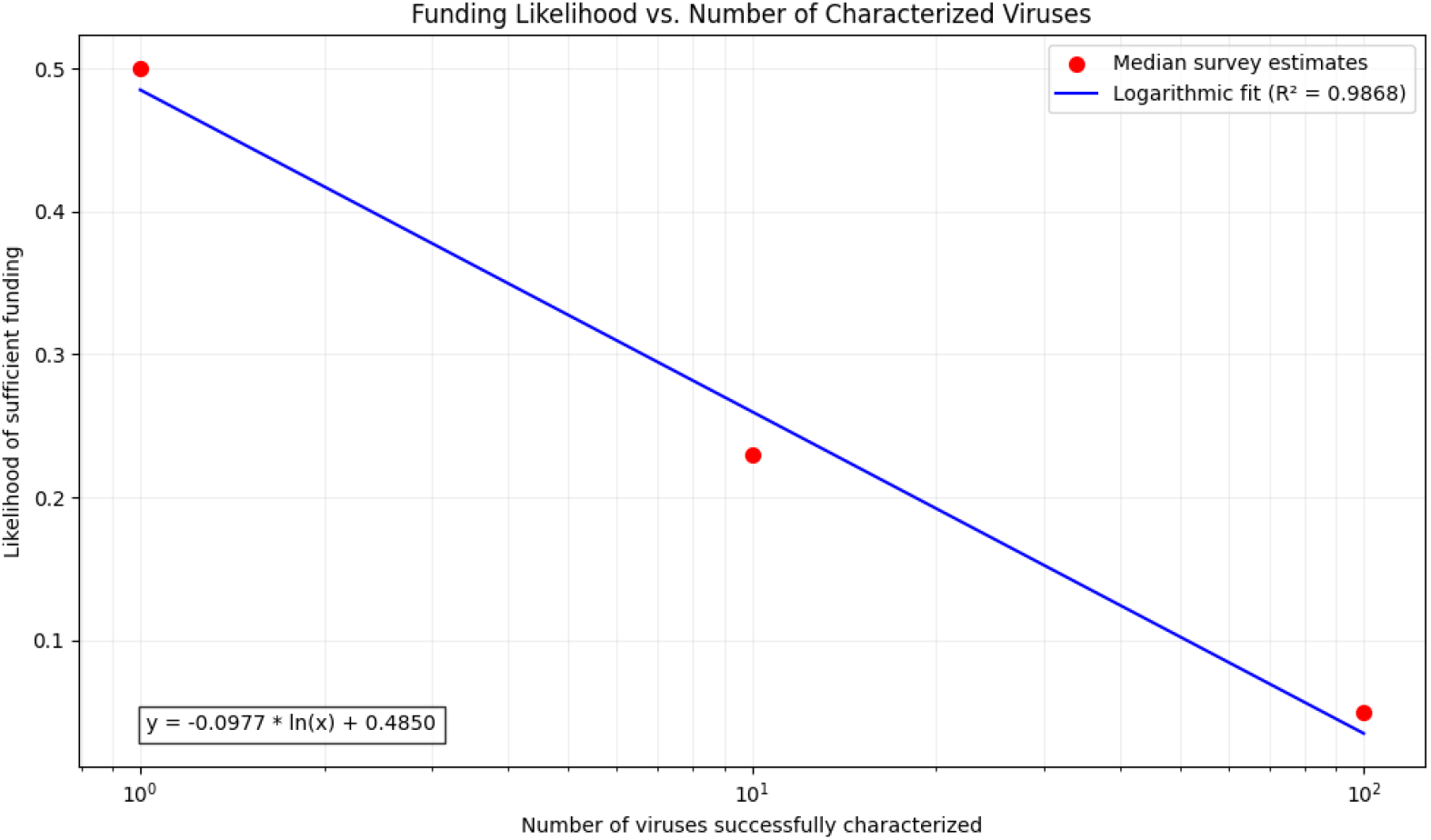
Relationship between the number of successfully identified pandemic-capable viruses and the likelihood of sufficient funding for countermeasure development. The line depicts a logarithmic curve fit to the data, y = -0.0977ln(x) + 0.485, where y is the likelihood of funding and x is the number of viruses.

In this scaled-up version of the PVI model, we assume the probabilities of funding for medical countermeasures and targeted non-pharmaceutical interventions are equal, such that there is a general probability targeted countermeasures will be funded *p*_*TCM*_, where *p*_*TCM*_ *= p*_*TMCM*_ = *p*_*NPI* | *PVI*_. We use the logarithmic curve generated by the survey data to generate estimates for *p*_*TCM*_ in the case where multiple pathogens are identified, such that:

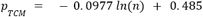

We plug this into equation (2), and plot the expected harm in scenarios where PVI and multiple pathogens are identified.

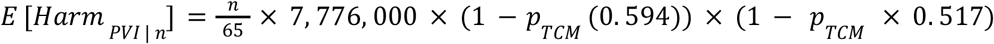

Consider the scenario where n = 2

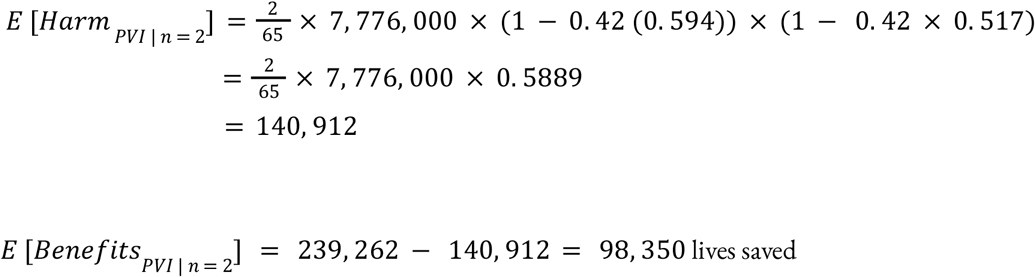

Using this approach, we estimated the lives saved in expectation based on the number of pandemic-capable viruses identified through PVI efforts. Given this, we estimate if all pandemic viruses are identified, this would save 642,000 lives in expectation, reducing overall natural pandemic risk by approximately 8% over the next decade.

### Uncertainty Quantification

Similar to the approach outlined in the VDS model, we assessed the uncertainty in the estimates for the benefits of identifying a single pandemic-capable virus and identifying all pandemic viruses. For the case of a single pandemic virus, we ran 100,000 MC simulations drawing parameter values directly from survey estimates for *p*_*TMCM*_, *p*_*NPI* | *PVI*_ and Δ*r*_*NPI* | *PVI*_ parameters. This resulted in a simulation mean and median of 48,911 and 47,725 lives saved, with a 90% central interval of 10,500 and 93,587 lives saved.

## Acknowledgements

We are grateful to Chris Said for help conducting the surveys, and to W. Bradshaw, M. Lipsitch, A. DeMarsh, S. Donoghue, N. Helm-Burger, O. Hershey-Wilson, and P. Medeiros for helpful comments. This work was supported by an Open Philanthropy Early-Career Award (to G.J.), NSF CAREER 1943141 (to K.M.E.), and by a gift from the Aphorism Foundation (to K.M.E.).

The authors declare no conflicts of interest.

