## Supplementary Information for "Modeling the benefits of virus discovery and pandemic virus identification"

### Survey Data

#### Initial Survey (n = 207)

1. Participant Research Areas and Affiliations
   *National Authorities* - 126
    - US CDC - 74
    - China CDC - 8
    - EU CDC (or member state equivalent) - 25
    - CDC equivalent - other countries - 52

   *Virus Detection & Characterization Programs* - 37
    - USAID Predict - 20
    - USAID DEEP VZN - 5
    - EcoHealth Alliance - 21
    - Global Virome Project - 12

   *Basic Research* - 118
    - Pathogens: Molecular or Cellular Biology - 90
    - Pathogens: Ecology or Evolution - 75

   *Applied Research* - 174
    - Vaccine Research - 54
    - Diagnostics Development - 78
    - Therapeutics Development - 38
    - Community Engagement and Training - 80
    - Policy or other preventative countermeasures - 108

2. Participant Role
    - Academia - 138
    - Industry - 6
    - Government - 70
    - Principal Investigator - 114
    - Staff Scientist or Postdoctoral fellow - 32
    - Graduate student -2
    - Predoctoral researcher/undergraduate student - 0

##### *Question 1 - Discovering Virus X: Effects on Broad-Spectrum Medical Countermeasures*

Question: How likely are we to have at least one approved broad-spectrum vaccine or therapeutic 10 years from now that will be effective against the next high-consequence pathogen (>1 million deaths)?

1 a) "Likelihood given our current knowledge of the global virome"
Parameter: $p_{BSV|\neg VDS}$

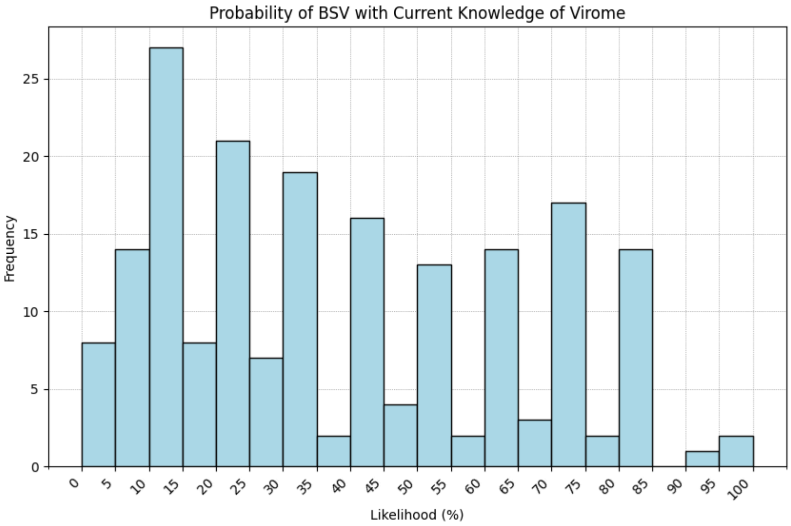

**Summary Statistics:**
Mean: 0.37 (90% CI [0.34, 0.4])
90% central interval: [0.05 , 0.8]
Distribution: Beta (0.96, 1.63)
St. dev: 0.26
Median: 0.3

1 b) "If we discovered and sequenced 3.0x as many viruses as today"
Parameter: $p_{BSV|3x\,dis}$

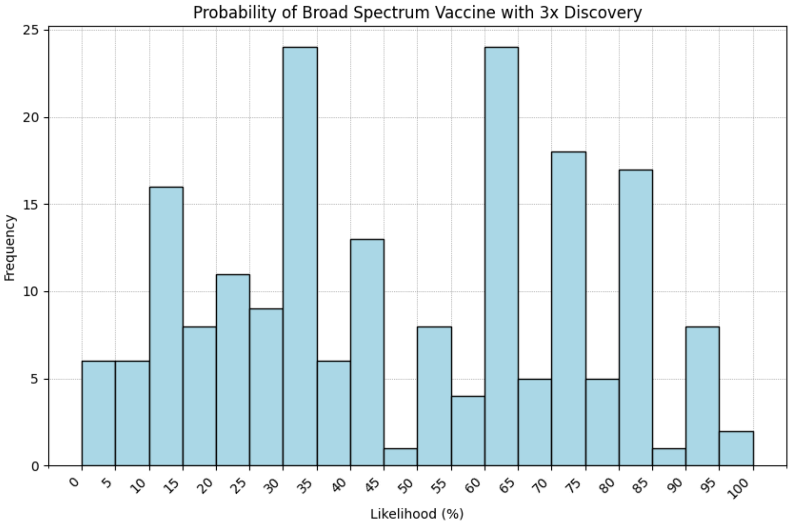

**Summary Statistics:**
Mean: 0.46 (90% CI [0.43, 0.49])
90% central interval: [0.06, 0.87]
Distribution: Beta (1.17, 1.40)
St.dev: 0.26
Median: 0.4

1c) "What if we discovered and sequenced all viruses in animals?"

Parameter: $p_{BSV\,|\,full\ virome}$

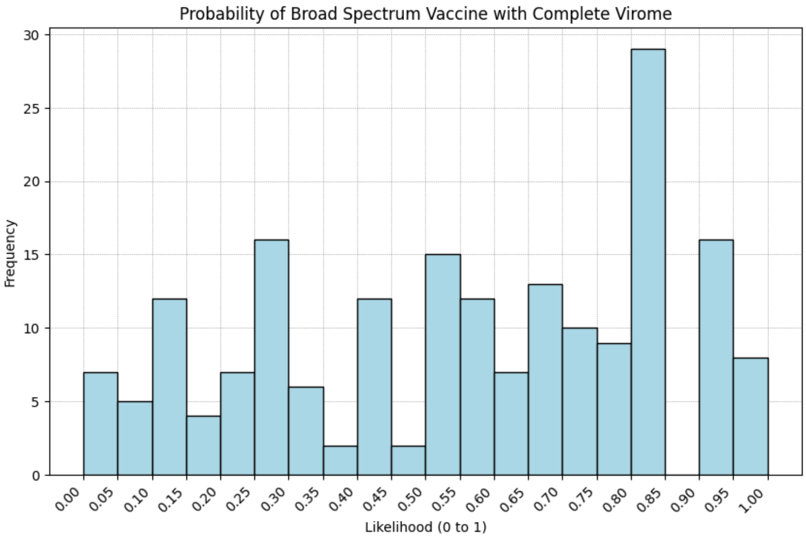

**Summary Statistics:**

Mean:0.55 (90% CI [0.51, 0.58])

90% central interval: [0.08, 0.95]

Distribution: Beta (1.14,0.93)

Median: 0.6

St. dev: 0.28

#### Change in Probabilities

Difference between 3x discovery and baseline (current knowledge of virome)

Parameter: $\Delta\, p_{BSV\,|\,VDS}$

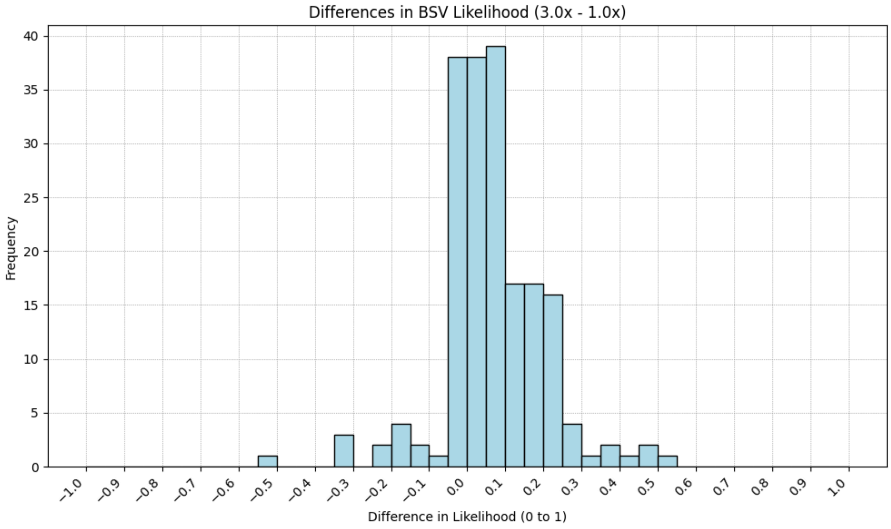

**Summary Statistics:**

Mean:0.08 (90%CI [0.07, 0.1]

90% central interval: [-0.13, 0.29]

Median: 0.09

St. dev: 0.13

Difference between complete virome and baseline (current knowledge of virome):

Parameter: $\Delta\, p_{BSV\,|\,\text{full}\ VDS}$

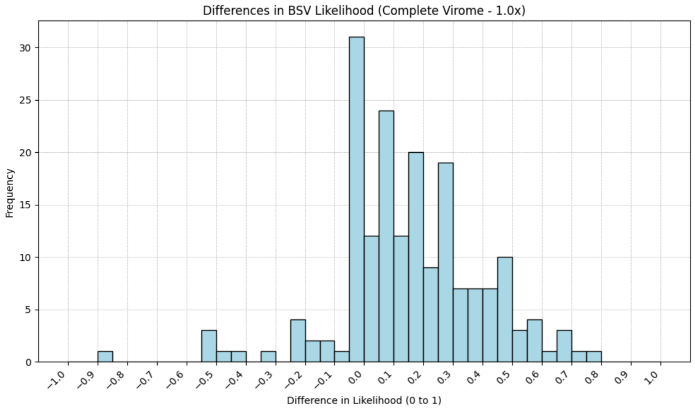

**Summary Statistics:**

Mean: 0.17 (90% CI [0.14, 0.19])

90% central interval: [-0.2, 0.58]

Median: 0.09

St. dev: 0.13

***Question 2 - Discovering Virus X: Effects on Preventative (Non-Medical) Interventions***

Suppose we pursue virus discovery and use this information to improve our broad-spectrum non-medical interventions.

*3x Discovery*

Question: " If we sequenced 3.0x as many viruses as today, estimate the relative reduction in pandemic risk from Virus X over the next 10 years relative to a world with no additional virus discovery (almost fully effective = "99%", no help at all = "0%")."

Parameter: $\Delta p_{NPI\,|\,VDS}$

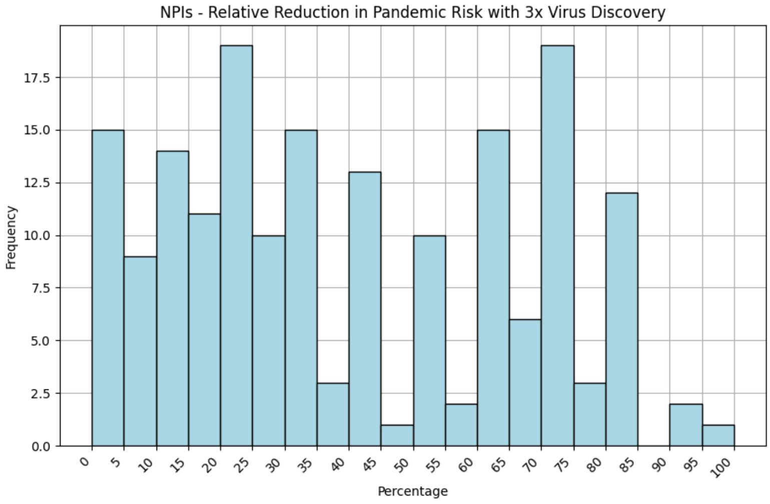

**Summary Statistics:**

Mean: 0.38 (90% CI [0.35,0.41])

90% central interval: [0.03, 0.86]

Distribution: Beta (0.95, 1.52)

St.dev: 0.26

Median: 0.3

IQR: 0.43

*Complete Virome*
2b) "What if we discovered and sequenced all viruses in animals?"

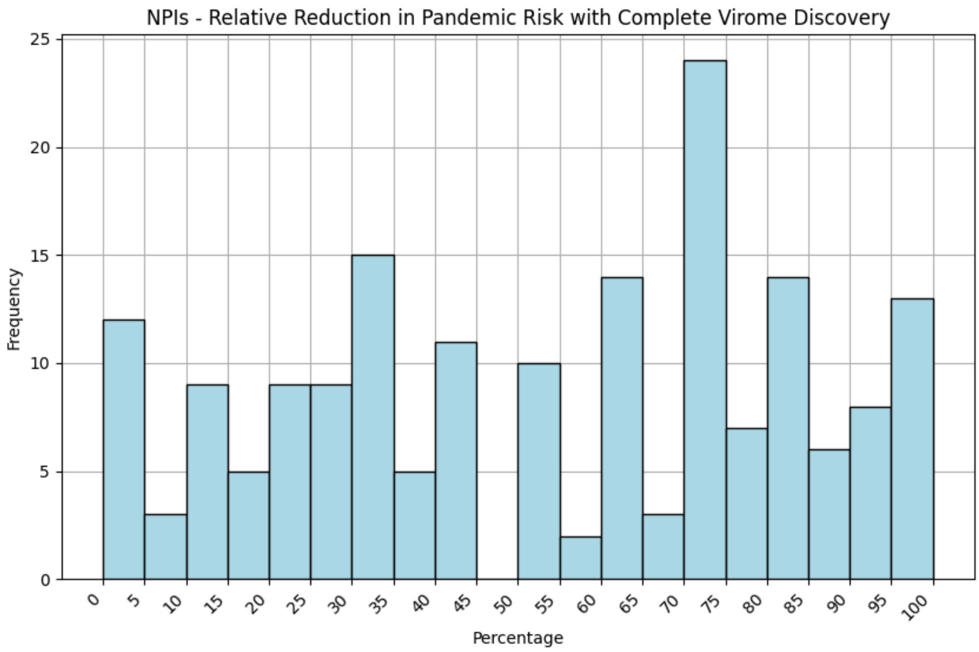

**Summary Statistics:**
Mean: 0.52 (90% CI [0.48, 0.55])
90% central interval: [0.05, 0.95]
Distribution: Beta(0.97,0.91)
Median: 0.57
St.dev: 0.3
IQR: 0.46

***Question 3 - High-Consequence Pathogens Circulating in Animal Reservoirs***
Question: "How many distinct viruses capable of sustained human-to-human transmission, with the potential to cause at least 1 million deaths, do you estimate are currently circulating in animal reservoirs around the world?"

- Less than 10 viruses
- 10 - 30 viruses
- 30 - 100 viruses
- 100 - 300 viruses
- 300 - 1k viruses
- 1k - 3k viruses
- 3k - 10k viruses
- 10k - 30k viruses
- More than 30k viruses

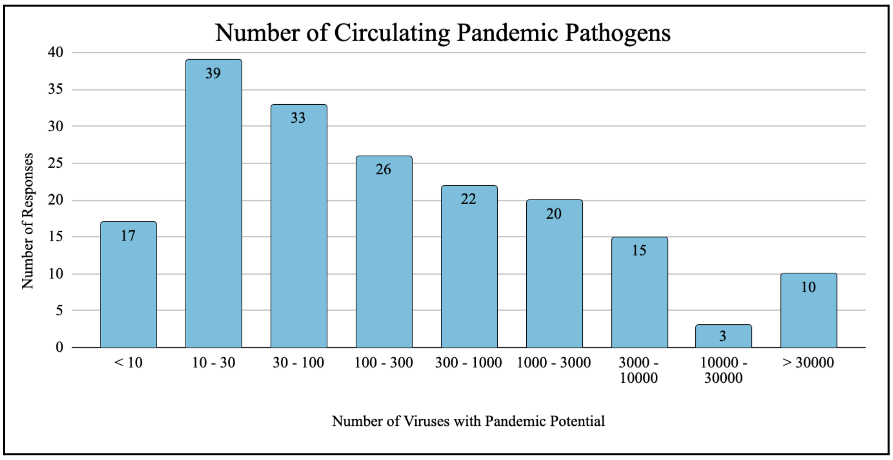

Summary Statistics:
Point estimate average: 172
90% central range: [6, 35250]
Distribution: LogN(5.1291,0.1924)

*90% CI calculated through bootstrapping method

***Question 4 - Pandemic Virus Identification: Effects on Targeted Medical Countermeasures***

Question: " If Virus X is characterized as pandemic-capable prior to spillover through PVI, what is the probability the world will invest enough funds to develop targeted MCMs before an outbreak begins?"

Parameter: $p_{TMCM}$

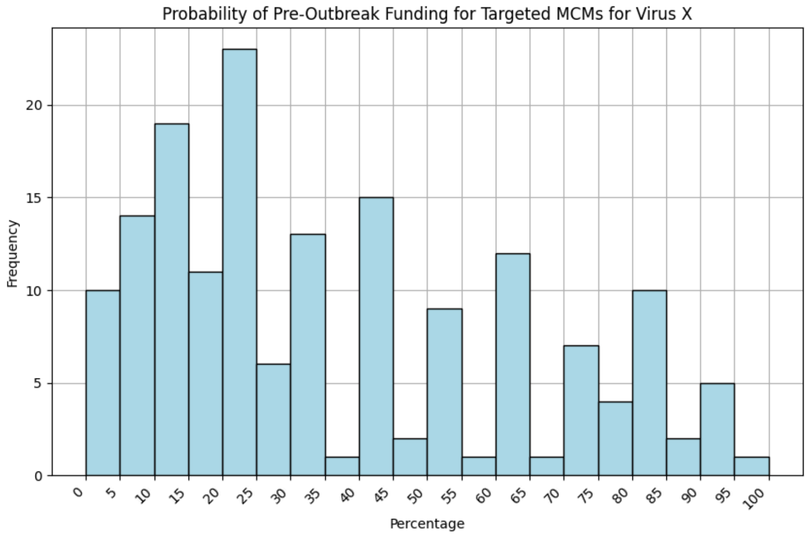

**Summary Statistics:**
Mean: 0.35 (90% CI [0.32,0.39])
90% central interval: [0.04, 0.83]
Median: 0.30
St.dev: 0.26
IQR: 0.47

***Question 5 - Acceleration of Medical Countermeasure Timelines due to PVI***

*Vaccines*
Question: "If the world does invest sufficient funds, a Virus X vaccine would be widely available (to at least 1 billion people) ____ days sooner relative to a world in which Virus X had not been characterized and flagged as a suspected pandemic risk in advance of the outbreak".

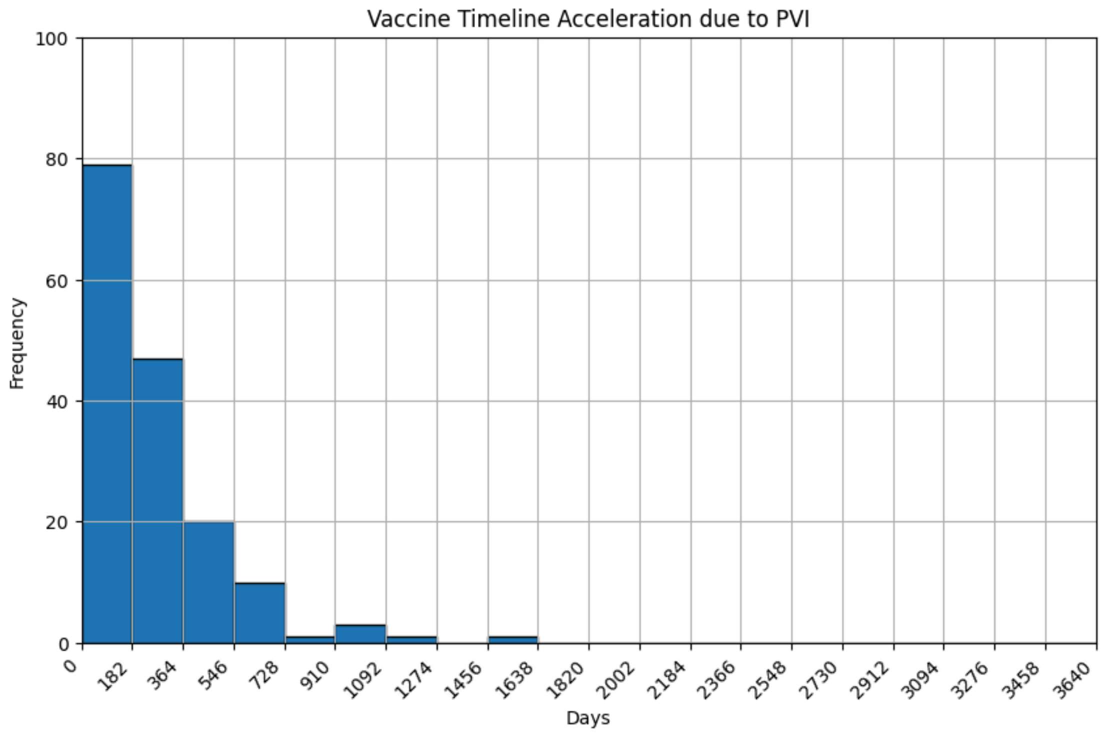

**Summary Statistics:**

Mean : 363 days

90% central interval: [7, 700]

St.dev:  1550 days

Median: 198 days (90% CI* [180, 200])

*used bootstrapping method for CI

*Therapeutics*

Question: "If the world does invest sufficient funds, a Virus X therapeutic would be widely available ____ days sooner, relative to a world in which Virus X had not been characterized and flagged as a suspected pandemic risk in advance of the outbreak".

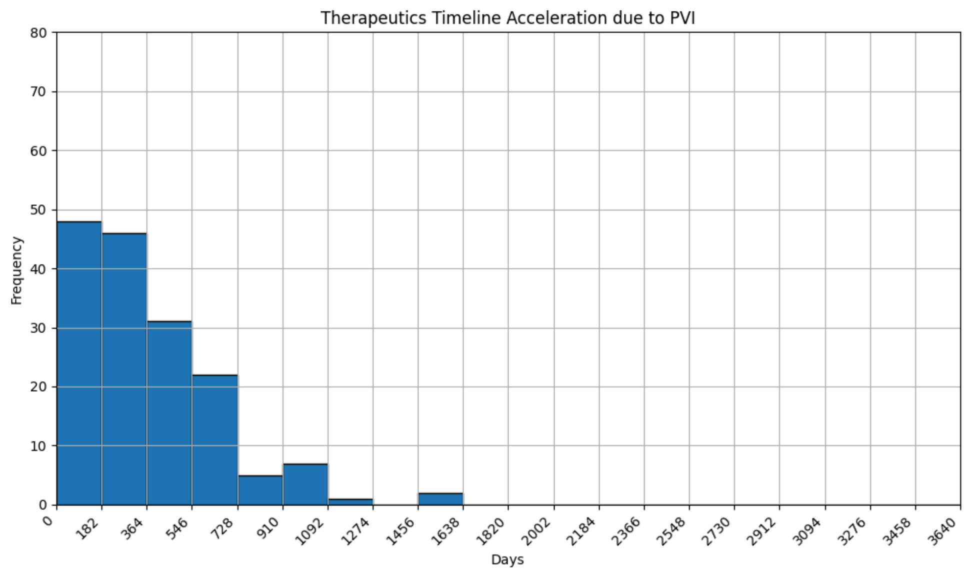

**Summary Statistics:**
Mean: 421 days
90% central interval: [10,995]
St.dev: 807 days
Median: 300 days (90% CI* [250, 360])
*used bootstrapping method for CI

##### ***Question 6 - Pandemic Virus Identification: Effects on Non-Medical Interventions***

Question: "How much would non-medical countermeasures targeting a characterized Virus X reduce the risk of a sustained outbreak over 10 years? Please input the reduction in risk relative to a world where Virus X is not characterized (almost fully effective = "99%", won't help at all = "0%")

Parameter: $\Delta r_{NPI\,|\,PVI}$

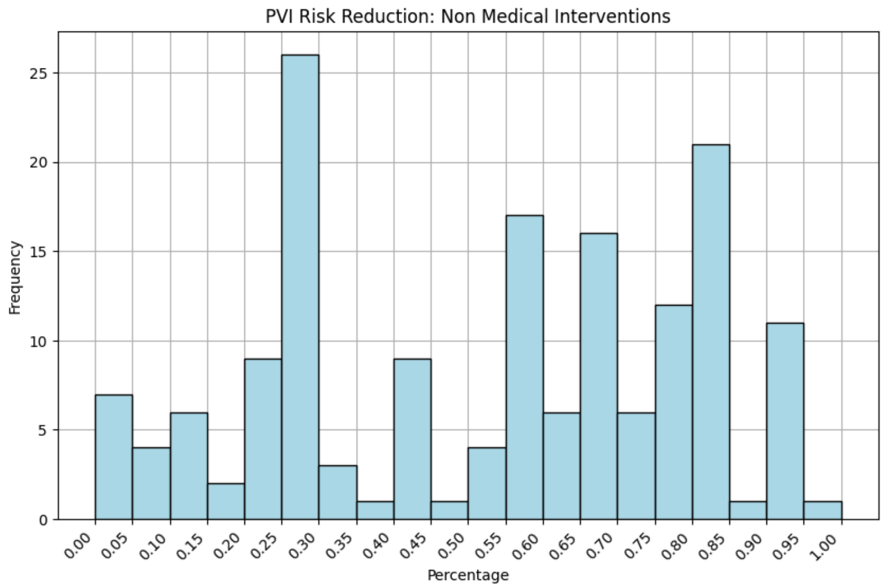

**Summary Statistics:**
Mean: 0.52 (90% CI [0.49, 0.56])
90% central interval: [0.09, 0.90]
Distribution: Beta (1.30,1.9)
St.dev: 0.27
Median: 0.6
IQR: 0.45

x̄: mean, M: median

| | National Academies | Virus Discovery and Characterization Programs | Basic Research | Applied Research |
|---|---|---|---|---|
| $\Delta p_{BSV \mid VDS}$ | x̄ = 0.08, M = 0.07 | x̄ = 0.13, M = 0.10 | x̄ = 0.10, M = 0.09 | x̄ = 0.08, M = 0.07 |
| $\Delta r_{NPI \mid VDS}$ | x̄ = 0.37, M = 0.3 | x̄ = 0.43, M = 0.4 | x̄ = 0.4, M = 0.38 | x̄ = 0.38, M = 0.3 |
| $\Delta p_{TMCM}$ | x̄ = 0.35, M = 0.29 | x̄ = 0.43, M = 0.4 | x̄ = 0.36, M = 0.3 | x̄ = 0.35, M = 0.29 |
| $\Delta r_{NPI \mid PVI}$ | x̄ = 0.53, M = 0.6 | x̄ = 0.54, M = 0.61 | x̄ = 0.54, M = 0.6 | x̄ = 0.53, M = 0.6 |
| Vaccine Timeline Acceleration | x̄ = 240, M = 180 | x̄ = 224, M = 180 | x̄ = 277, M = 200 | x̄ = 245, M = 196 |
| Therapeutics Timeline Acceleration | x̄ = 451, M = 300 | x̄ = 318, M = 205 | x̄ = 394, M = 300 | x̄ = 427, M = 300 |

#### Follow-up Survey (n = 42)

***Question 1 - Risk Distribution***

Question: “Which of the following best match with your assessment of how pandemic risk is distributed across potential high-consequence pathogens? Note that all percentages below are as a fraction of the potentially high-consequence pathogens only; pathogens without pandemic potential are not included here.

1. All pathogens in this category are equally risky: the top 20% most risky pathogens contribute 20-30% of expected mortality
2. Narrow-tailed distribution: there is some difference between pathogens, but only a minority of the risk is concentrated in the 20% most risky pathogens (30-50% of expected mortality)
3. Fat-tailed distribution: most, but not all, of the risk is concentrated in the top 20% most risky pathogens (50-90% of expected mortality)
4. Very fat-tailed distribution: nearly all risk is concentrated in the top 20% most risky pathogens (defined as 90-100% of expected mortality)”

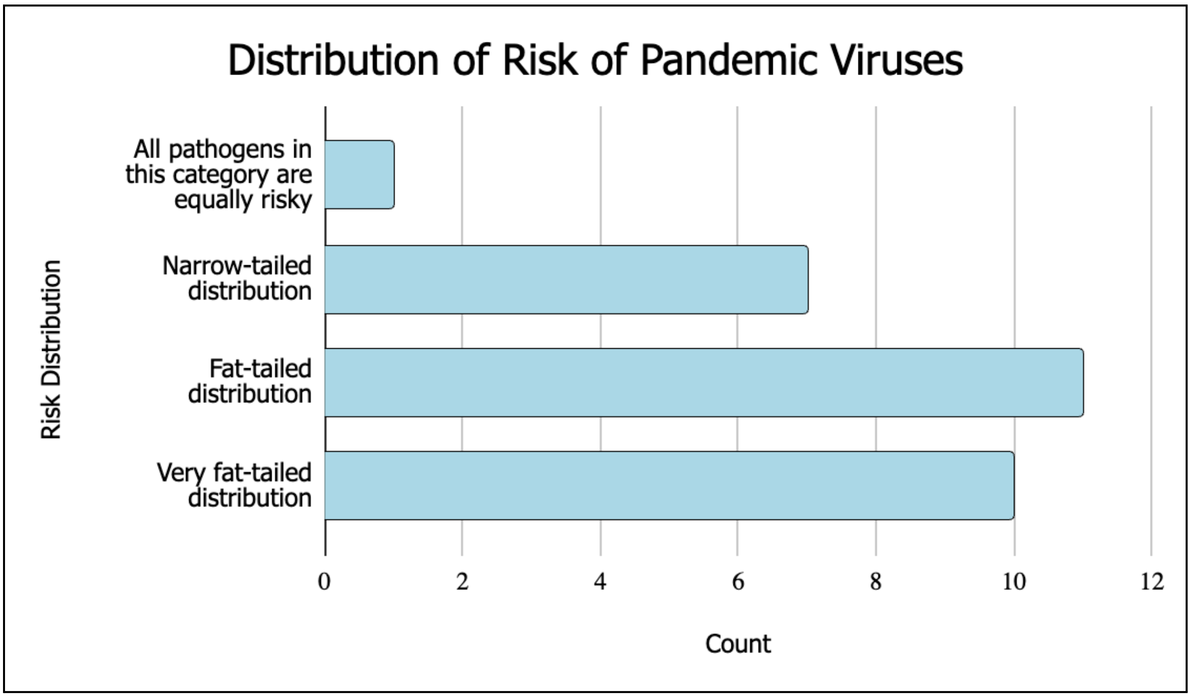

Weighted Average of Risk Distribution Results:

Weighted average was calculated using the midpoints of the expected mortality ranges and the number of respondents as weights.

$$\text{Weighted Average} = \frac{(9.5 + 7.7 + 2.8 + 0.25)}{29} = 0.70$$

The top 20% of most risky pathogens contribute to 70% of the expected mortality.

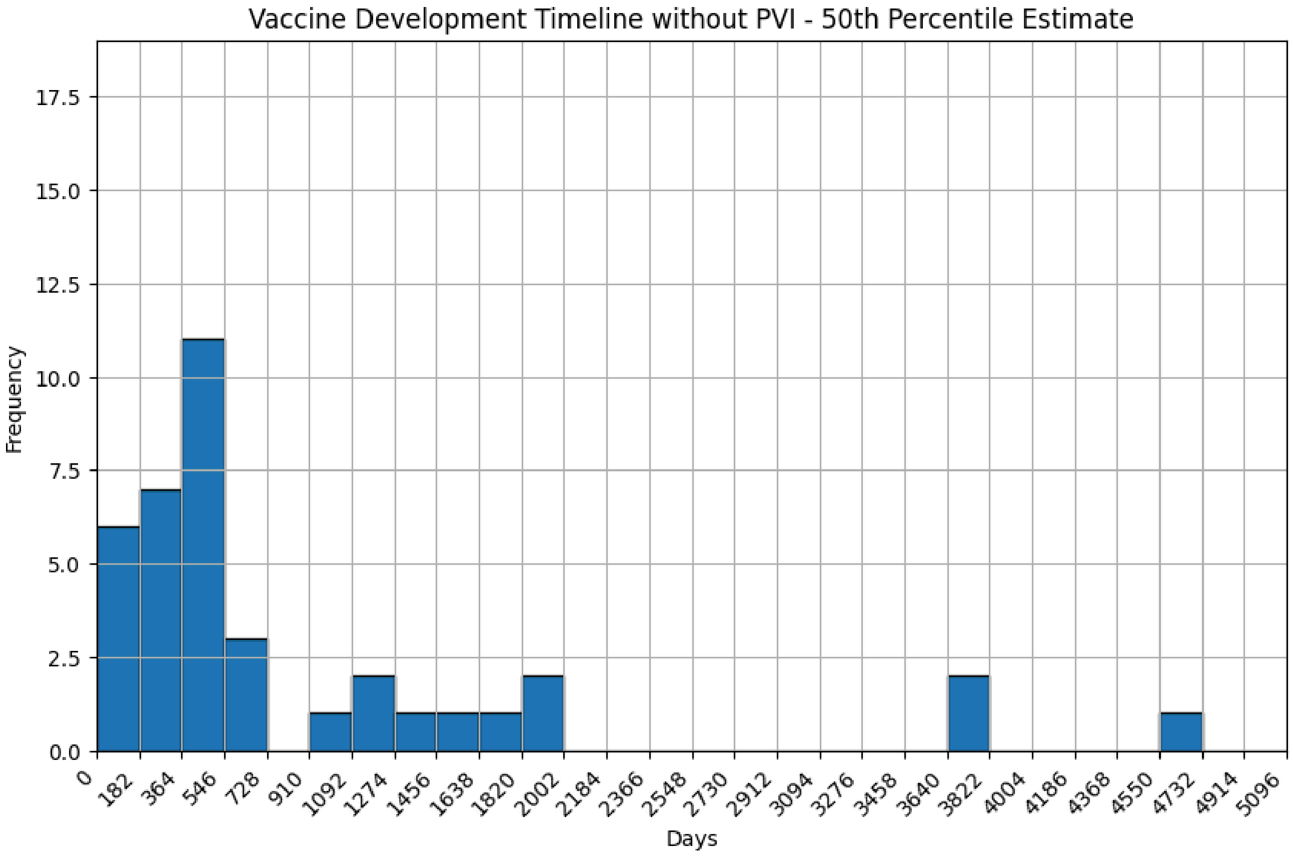

***Question 2a - Vaccine Development without PVI***

Summary Statistics:

Mean: 858 days

St.dev: 1050 days

Median: 382 days

IQR: 756 days

*90K estimate outlier removed

***Question 2b - Vaccine Development with PVI***

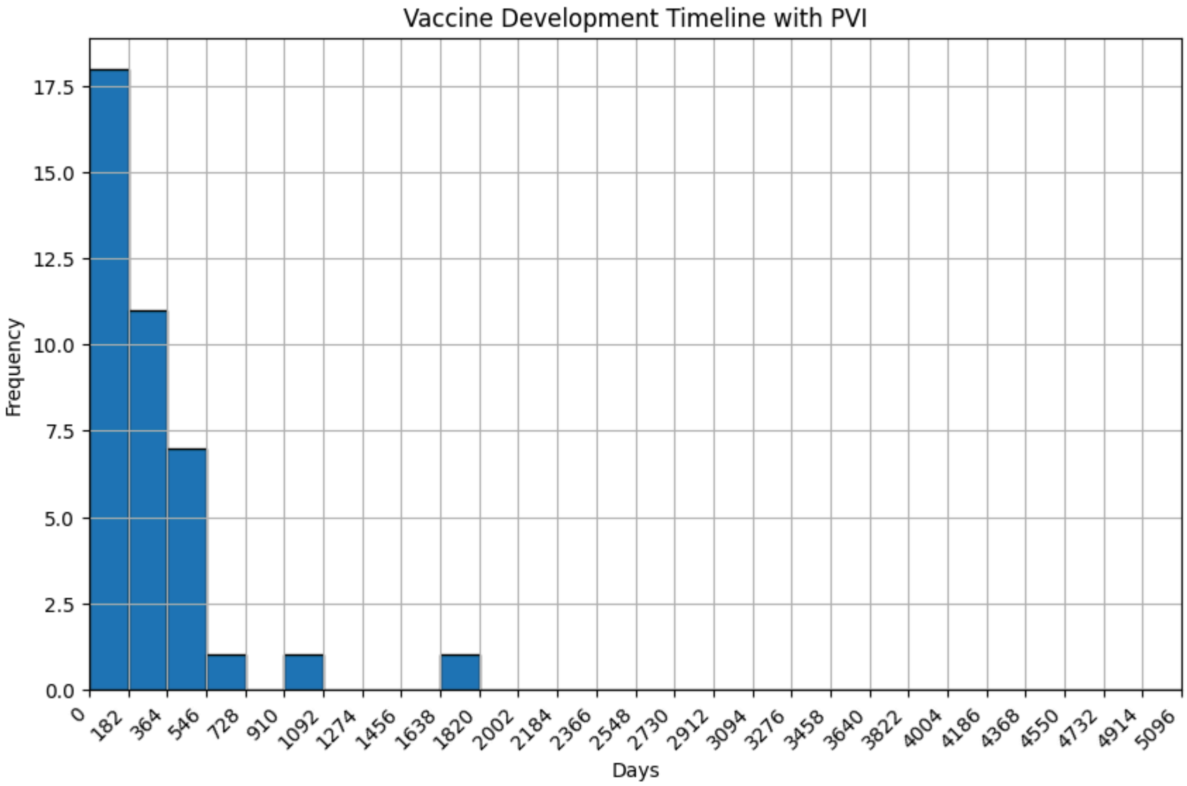

Summary Statistics:
Mean: 287 days
St.dev: 303 days
Median: 200 days
IQR: 213 days

Note: Median difference reported in followup survey regarding timelines

***Question 3 - Funding***

Parameter: $p_{TCM}$

1. If only one (1) virus is identified in the laboratory as a suspected pandemic threat over the next ten years, what is the likelihood that the world invests enough funds to develop targeted countermeasures against this virus?
2. If ten (10) different viruses are identified in the laboratory as suspected pandemic threats over the next ten years, what is the likelihood that the world invests enough funds to develop targeted countermeasures against at least one (1) of these viruses?

3. If ten (10) different viruses are identified in the laboratory as suspected pandemic threats over the next ten years, what is the likelihood that the world invests enough funds to develop targeted countermeasures against all ten (10) viruses?
4. "If a hundred (100) different viruses are identified in the laboratory as suspected pandemic threats over the next ten years, what is the likelihood that the world invests enough funds to develop targeted countermeasures against at least one (1) of these viruses?
5. If a hundred (100) different viruses are identified in the laboratory as suspected pandemic threats over the next ten years, what is the likelihood that the world invests enough funds to develop targeted countermeasures against all one hundred (100) viruses?

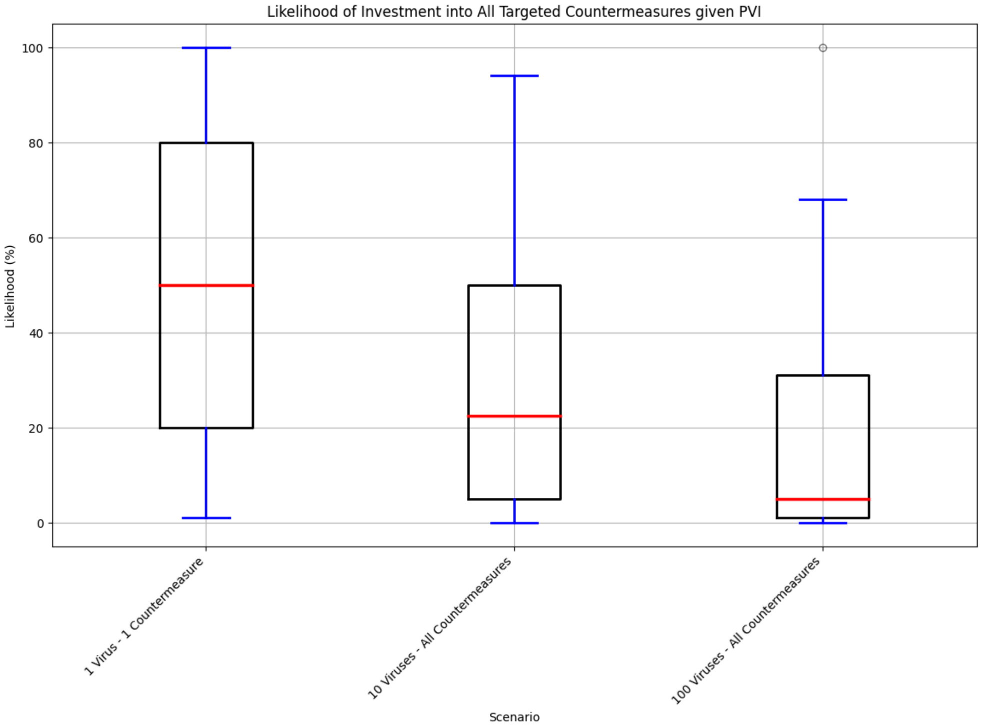

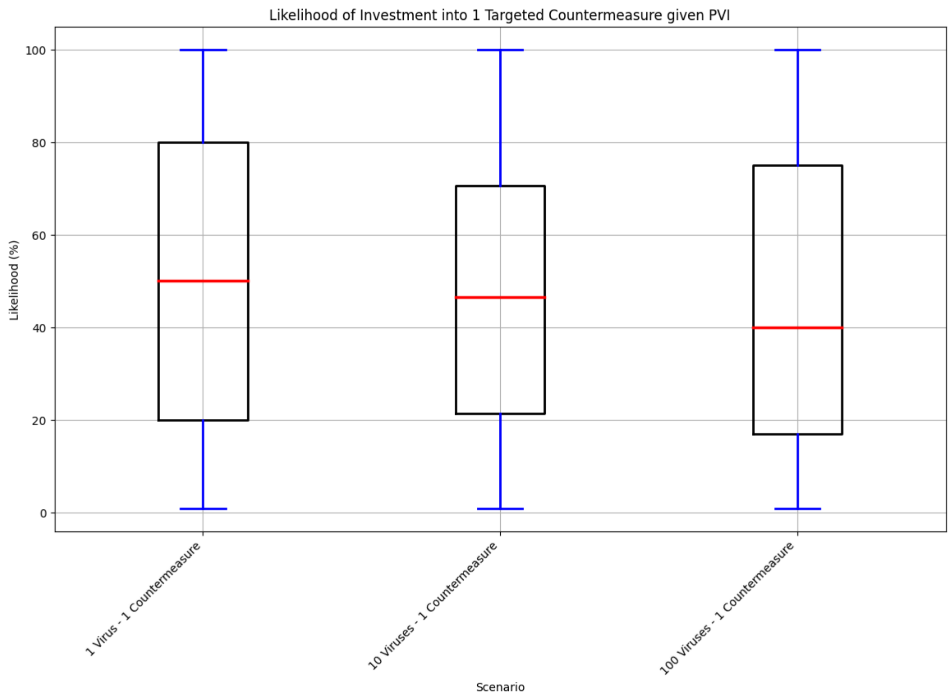

**Summary Statistics**

| | **1 Virus - 1 MCM (%)** | **10 Viruses - 10 MCMs (%)** | **100 Viruses - 100 MCMs (%)** |
|---|---|---|---|
| Mean | 0.48 | 0.32 | 0.18 |
| 90% central interval | (0.02, 0.94) | (0.01, 0.81) | (0.0, 0.55) |
| St.dev | 0.32 | 0.29 | 0.25 |
| Median | 0.5 | 0.2 | 0.05 |
| Distribution | Beta (0.72,0.84) | Beta (0.53,1.18) | Beta (0.27,1.20) |

##### *Question 4 - PVI*

Question: "Considering only the risk of a pandemic of zoonotic origin, do you think pandemic characterization on net decreases the risk from Virus X, increases the risk, or does not change the level of risk?

A. Pandemic characterization decreases the risk from pandemics of zoonotic origin, on net

B. Pandemic characterization does not make a difference to the risk from pandemics of zoonotic origin
C. Pandemic characterization increases the risk from pandemics of zoonotic origin, on net
"

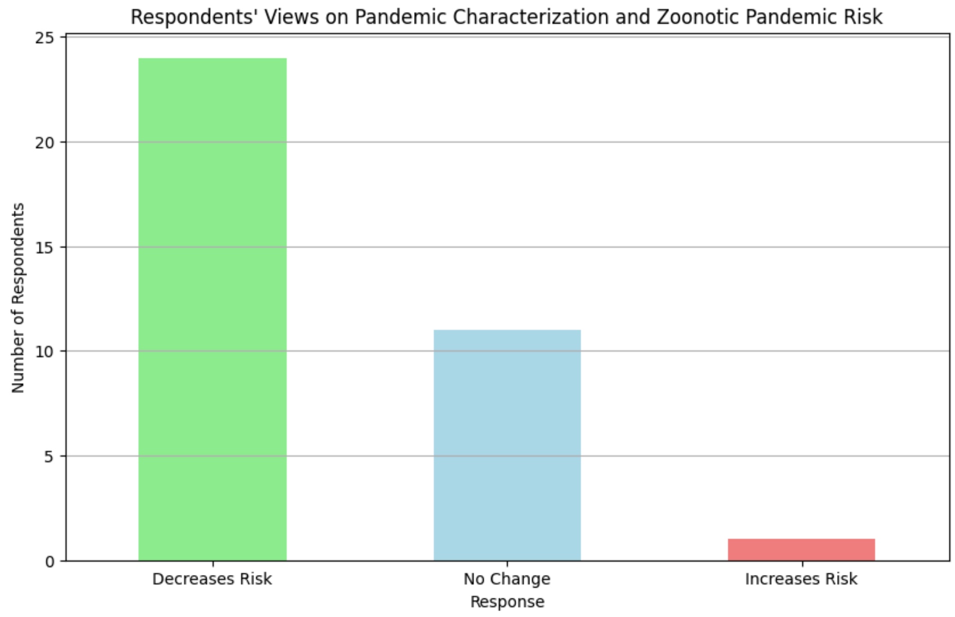

Decreases Risk: 24
No Change: 11
Increases Risk: 1

#### Model Parameters

The table below presents all the parameters of our model, noting the parameter name, point estimate, distribution for parameters estimated with survey data, as well as the description and source of the data for each parameter.

Table S1: Model parameters and relevant information

| **Parameter** | **Median/Central estimate, [90% Central Range]** | **Description** | **Source** |
|---|---|---|---|
| $E[Harm_{VDS}]$ | 96,901 deaths [0, 1.46 million] | Expected number of deaths from natural pandemics per virus per decade given viral discovery efforts triple the number of viruses we know about | Model |

| | | | |
|---|---|---|---|
| $E\,[Harm_{PVI}]$ | 48,000 deaths [10,500 ; 93,600] | Expected number of deaths from natural pandemics per virus per decade given we characterize the pandemic potential of the virus | Model |
| $v_{total}$ | 172 [6, 35250] | Total number of pandemic-capable viruses circulating in the world | Survey |
| $p_{BSV\,\vert\,\neg VDS}$ | 30% [5% , 80%] | Baseline probability at least one approved broad-spectrum vaccine for use against Virus X prior to spillover without any additional VDS | Survey |
| $p_{BSV\,\vert\,3x\,dis}$ | 40% [6%, 87%] | Probability discovering 3 times as many viruses as today results in at least one approved broad-spectrum vaccine for use against Virus X prior to spillover | Survey |
| $p_{BSV\,\vert\,full\ virome}$ | 60% [8%, 95%] | Probability discovering all viruses in animals results in at least one approved broad-spectrum vaccine for use against Virus X prior to spillover | Survey |
| $\Delta\, p_{BSV\,\vert\,VDS}$ | 9% [-13%, 29%] | Relative increase in likelihood of an approved broad-spectrum vaccine prior to Virus X spillover given 3x virus discovery | $p_{BSV\,\vert\,3x\,dis} - p_{BSV\,\vert\,\neg VDS}$ |
| $\Delta\, p_{BSV\,\vert\,full\ VDS}$ | 17.9% [-13%, 29%] | Relative increase in likelihood of an approved broad-spectrum vaccine prior to Virus X spillover given complete virome discovery | $p_{BSV\,\vert\,full\ VDS} - p_{BSV\,\vert\,\neg VDS}$ |
| $\Delta p_{NPI\,\vert\,VDS}$ | 14% [0%, 56%] | Relative change in probability non-pharmaceutical interventions would be directed towards Virus X hotspot due to 3x viral discovery | Survey, inferred |
| $\Delta p_{NPI\,\vert\,full\ VDS}$ | 25% [0%, 87%] | Relative change in probability non-pharmaceutical interventions would be directed towards Virus X hotspot due to complete virome discovery | Survey, inferred |
| $m_{BSV}$ | 41.6% | reduction in harm due to the availability of a broad-spectrum vaccine for Virus X prior to spillover | Literature |
| $\Delta r_{NPI\,\vert\,VDS}$ | 30% [0.03, 0.86] | Relative reduction in harm due to increased targeted of non-pharmaceutical interventions towards Virus X hotspot due to 3x viral discovery | Survey |
| $\Delta r_{NPI\,\vert\,full\ VDS}$ | 57% [5%, 96%] | Relative reduction in harm due to increased targeted of non-pharmaceutical interventions towards Virus X hotspot due to complete virome | Survey |
| $p_{TMCM}$ | 30% [4%, 83%] | Probability of targeted Virus X medical countermeasures receiving sufficient funds for development prior to spillover and outbreak | Survey |
| $p_{NPI\,\vert\,PVI}$ | 48% [2%, 94%] | Probability PVI results in targeted non-pharmaceutical interventions at the Virus X hotspot | Survey |
| $\Delta m_{TV}$ | 54% [2%, *] | Relative reduction in harm due to earlier release of targeted vaccines | Literature + Survey |

| | | | |
|---|---|---|---|
| $\Delta m_{TT}$ | 9% [0.4%, *] | Relative reduction in harm due to earlier release of targeted therapeutics | Literature + Survey |
| $\Delta r_{NPI \mid PVI}$ | 60% [9%, 90%] | Relative reduction in risk due to improved non-pharmaceutical interventions from PVI | Survey |

* was used to denote where the upper bound estimate was greater than 100% due to participants' upper bound estimates of accelerated vaccine and therapeutics development timelines, which would have resulted in more lives saved than those lost in pandemic events.

### Parameter Estimation

#### Number of Pandemic-Capable Viruses

In our model, we used survey data to estimate (1) the total number of pandemic-capable viruses currently circulating around the world, $v_{total}$; and (2) the statistical equivalent number of viruses with equally likely probabilities of seeding a pandemic event, $v$.

To estimate (1) **the total number** we applied a log transformation to the lower and upper bound of each category in question 3 of the initial survey and took the log midpoint, using 1 as the lower bound for the "less than 10 viruses" category, and 100,000 as the upper bound for the "more than 30000 viruses" category. We then used these midpoints to calculate the weighted average., using the number of respondents for each category as weights. We then exponentiated this weighted average to convert the value back to the normal scale.

$$Weighted\ log\ average\ = \frac{(0.5 \times 17 + 1.24 \times 39 + 1.74 \times 33 + 2.24 \times 26 + 2.74 \times 22 + 3.24 \times 20 + 3.74 \times 15 + 4.24 \times 3 + 4.74 \times 10)}{185}$$

$$Weighted\ log\ average\ = \ 2.23$$

$$Average\ = \ 10^{2.23}\ = \ 172$$

The median category was 30-100 viruses. Using a similar approach, this would result in a median of $10^{1.74}$ = 55 viruses.

However PVI efforts are most likely to identify those circulating at high-risk hotspots which are at greatest risk of spilling over in human populations, as viral discovery efforts tend to target key taxa that are most likely to carry zoonotic viruses, such as non-human primates. Participants noted the top 20% of most risky pathogens contribute to 70% of the expected mortality in the followup survey. This difference in risk can be explained by differing environmental factors, where some viruses are in locations with high risk of zoonotic spillover to humans, while others are likely to be circulating in hosts that have little contact with humans. To address this diversity in likelihood in our calculations, we adjusted the 172 estimate using this risk distribution to get the (2) **effective number of pandemic capable viruses** with equally likely probabilities of seeding a pandemic event. This estimate is based on the assumption that the likelihood of identifying a pandemic-capable pathogen is proportional to its contribution to pandemic risk. On net, we expect it generates a favorable (higher) estimate of the potential benefits from PVI, since some of the variation in risk from the pandemic-capable pathogens may be unrelated to the likelihood of identifying it, which would increase the effective number of pandemic-capable viruses.

20% of pathogens contribute to 70% of the risk → $0.2 \times 172 = 34$
80% of pathogens contribute to the remaining 30% of the risk → $0.8 \times 172 = 138$

Let $v$ be the *effective number of pandemic-capable viruses*. We use a weighted approach, using the proportion of risk each group of pathogens contributes to as the weights.

$$0.7 \times \frac{34}{v} + 0.3 \times \frac{138}{v} = 1$$
$$v = 34 \times 0.7 + 138 \times 0.3$$
$$v = 65.2 \approx 65$$

This means that if a single pathogen is identified as pandemic-capable, in expectation it is linked to 1/65th of the aggregate zoonotic pathogen risk.

#### Vaccines

Both VDS and PVI have the potential to influence timelines associated with vaccine development. We establish the baseline Virus X scenario as one where there is initially no vaccine available, and a targeted vaccine becomes available 357 days after the outbreak begins.

With VDS, we consider the scenario where a **broad-spectrum** vaccine (either pan-genus or pan-species) is developed and approved prior to the Virus X outbreak, with manufacturing and distribution immediately scaled up as soon as the outbreak begins. We do not assume that a large fraction of humanity (>500m) will be vaccinated at pandemic onset since even seasonal influenza vaccines fail to reach such an uptake level.

With PVI, we consider the scenario where a **targeted** vaccine begins development prior to the outbreak rather than after the virus has spilled over, resulting in an accelerated vaccine approval and distribution. The additional deaths averted come from this earlier release.

To generate quantitative estimates, we use COVID-19 data to both establish a baseline as well as evaluate the alternate scenarios. The figure below illustrates Moderna's COVID-19 vaccine development timeline, as well as the duration it took to vaccinate 1 billion people across all manufacturers.

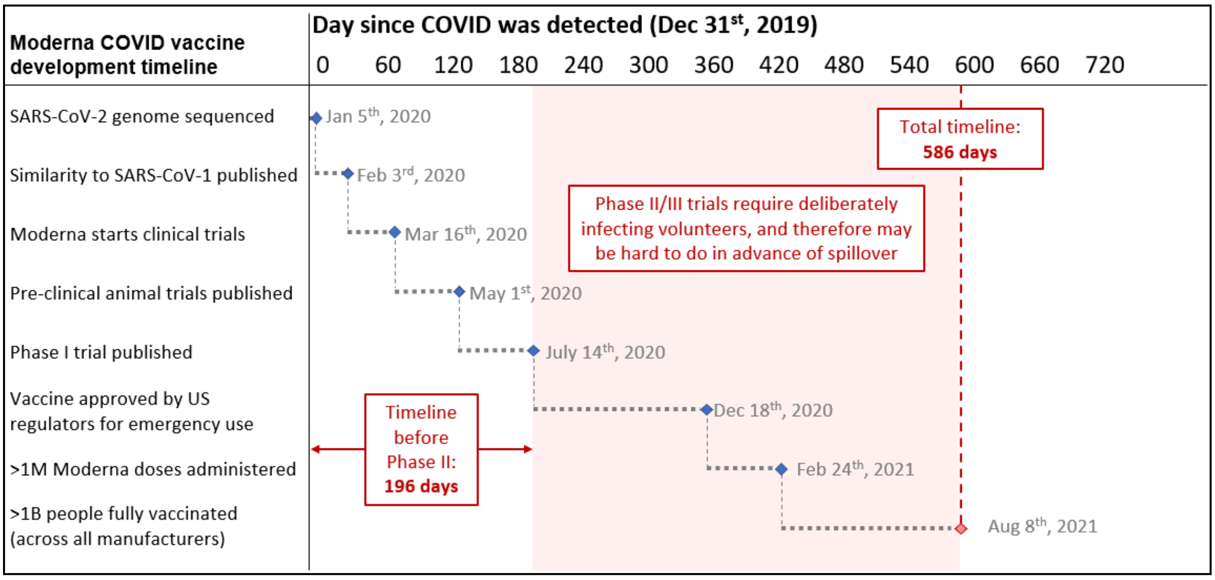

**Supplementary Figure 1 | Vaccine development timeline for COVID-19.**

##### *Broad-Spectrum Vaccines* ($m_{BSV}$)

The parameter $m_{BSV}$ in the model represents the reduction in harm due to the release of a broad-spectrum release at the beginning of the pandemic. We estimate this parameter by using data from the first two years of the COVID-19 pandemic to establish the baseline cumulative death toll and vaccine coverage. In this scenario, we make the following assumptions:

- The broad-spectrum vaccine is released 300 days prior to the release of the targeted vaccine, approximately 50 days into the pandemic.
- Vaccine distribution follows the same trajectory as the targeted COVID-19 vaccines in the U.S. and U.K.
- The broad-spectrum vaccines are half as effective as targeted vaccines at preventing mortality.

Więcek et al. estimate 240,715 additional lives could have been saved between the U.S. and U.K. if the COVID-19 vaccine was released 90 days earlier. We extrapolate that if the vaccine had been released 300 days earlier (approximately 50 days into the outbreak), this would have resulted in an additional 802,383 lives saved between the U.S. and U.K. Using our assumption that a BSV would be half as effective as a targeted vaccine, a BSV being released 300 days prior to the release of a targeted vaccine would result in an additional 401,191 lives saved within the U.S and U.K. Given the U.S. and U.K. reported a total of 964,000 COVID-19 deaths within the first two years, we estimate mBSV to be:

$$m_{BSV} = \frac{401,191}{964,000} = 0.416$$

##### *Accelerated Targeted Vaccines* ($\Delta m_{TV}$)

The parameter $\Delta m_{TV}$ represents the additional reduction in harm due to the earlier release of a targeted vaccine. In our expert survey, the median response of how much earlier a targeted vaccine would be released was 198 days. In our followup survey, when asked how long it would take to develop a targeted vaccine without PVI, participants provided a median estimate of 382 days, closely matching the 357 days it took to from the beginning of the COVID-19 pandemic to when the Pfizer-BioNTech vaccine received emergency use approval. Similar to above, we estimate $m_{TV}$ by considering the counterfactual scenario where the targeted COVID-19 mRNA vaccines were approved for use 198 days earlier. To estimate the additional deaths prevented, we primarily draw from the 2023 Więcek et al. study which estimated the potential lives saved by earlier COVID-19 vaccination in scenarios where vaccines were available 30, 60, or 90 days earlier than the actual timeline.

They estimate that within the first two years (by Jan 2022), between the U.S. and U.K. 240,715 [117,731; 332,397] additional deaths would have been prevented if targeted vaccines were released 90 days sooner, for an average of 2675 deaths prevented per day between these two countries. Over the course of 182 days, this would result in 524,300 deaths prevented between the two countries. The U.S. and U.K. reported a total of 964,000 COVID-19 deaths by Jan 2022. This suggests roughly 0.54 an additional life could have been saved for every death recorded.

$$\Delta m_{TV} = \frac{524300}{964000} = 0.54$$

Given there were approximately 5.49 million deaths globally recorded by this time, we estimate that globally, this means a targeted vaccine released 198 days earlier would save an additional 2.965 million lives globally for a pandemic similar to COVID-19.

#### Therapeutics

##### *Accelerated Targeted Therapeutics* ($\Delta m_{TT}$)

The parameter $\Delta m_{TT}$ represents the additional reduction in harm that would come from an earlier release of a targeted Virus X therapeutic due to PVI efforts. To estimate $m_{TT}$ , we considered how much earlier the therapeutic would be released, how effective it would be, and how many of those infected would have access to the therapeutic. During the COVID-19 pandemic, Pfizer developed Paxlovid (nirmatrelvir–ritonavir) , an orally administered antiviral therapy. Paxlovid was approved for use 722 days into the outbreak, and was reported to reduce the risk of hospitalization or death by 88% amongst unvaccinated high-risk patients with COVID-19. We use these values as proxies for the efficacy and baseline timeline of a targeted Virus X therapeutic.

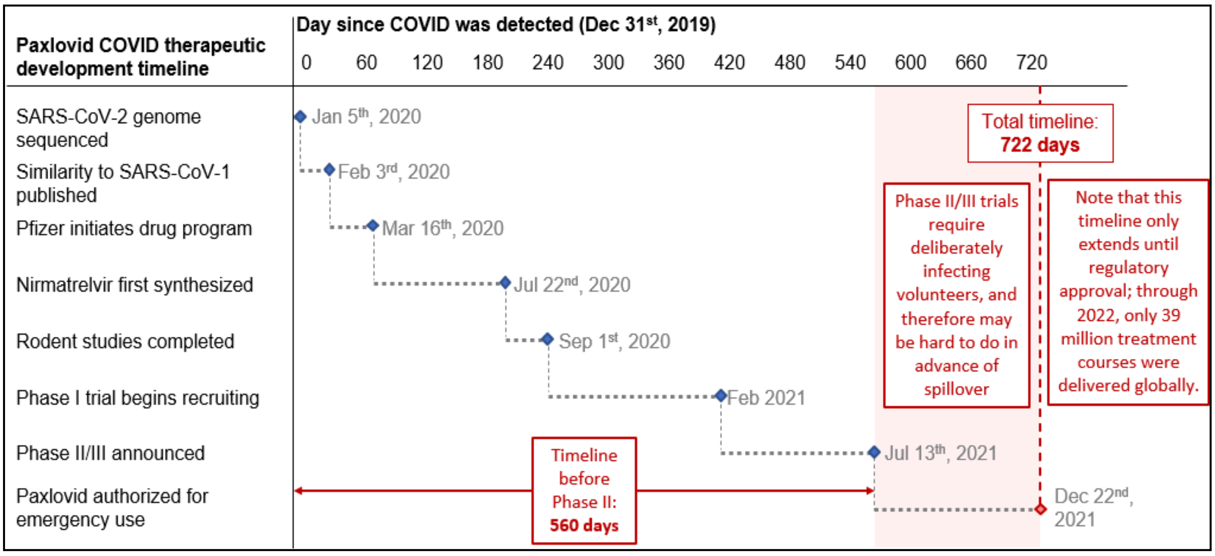

**Supplementary Figure 2 | Therapeutic development timeline for COVID-19.**

According to our survey, the median response to how many days sooner a Virus X therapeutic would be available if sufficient funds were invested and the virus had been characterized and flagged in advance was 300 days. We'll estimate $m_{TT}$ by estimating the additional reduction in harm that could have been achieved if Paxlovid was approved February 24, 2021. On May 5th 2023, the WHO declared the end of the COVID-19 pandemic, at which point 6.93 million deaths were recorded. We use this as the baseline number of Virus X pandemic deaths.

- $t_{base}$ = 722 days: Dec 22 2021 → 5.3M deaths
- $t_{PVI}$ = 422 days: Feb 22 2021 → 2.62M deaths

There were 2.68 million deaths between $t_{base}$ and $t_{PVI}$. The CDC reported a 28.4% adoption rate amongst eligible patients in the U.S. between April and August of 2022 (Shah et al. 2022). Using these values, we estimate that 650,157 additional lives could have been saved during that period through access to Paxlovid.

$$\begin{aligned} Additional\ Lives\ Saved &= deaths\ during\ 300\ days\ \times\ adoption\ rate\ \times\ efficacy \\ &= 2,680,000\ \times\ 0.282\ \times\ 0.88 \\ &= 650,179 \end{aligned}$$

Now taking into consideration the total deaths recorded during the pandemic:

$$\Delta m_{TT} = \frac{additional\ lives\ saved}{baseline\ pandemic\ deaths} = \frac{650{,}179}{6{,}930{,}000} = 0.094$$

Therefore we estimate that if a targeted therapeutic were to be accelerated by 300 days due to PVI efforts, this would result in an additional 9.4% of deaths prevented during a Virus X pandemic.

### Model Calculations

#### Virus Discovery

To evaluate the benefits of VDS, in our paper we define a scenario with VDS as a scenario in which 3 times as many viruses as today were discovered and sequenced. We also additionally evaluate the benefits of complete VDS, that is discovering and sequencing the entire virome, which can serve to establish an upper bound of the value of these efforts.

The key parameters in the VDS portion of this model are: the change in probability of a broad-spectrum vaccine being approved for use against Virus X, $\Delta p_{BSV|VDS}$; the conditional reduction in harm if a broad-spectrum vaccine (BSV) were available, $m_{BSV}$; the change in probability non-pharmaceutical interventions would be better targeted and directed towards Virus X hotspot(s) due to VDS, $\Delta p_{BSV|VDS}$; and the relative reduction in risk from better targeted non-pharmaceutical interventions, $\Delta r_{NPI\,|\,VDS}$.

In our survey we asked participants to estimate likelihood of having at least one approved broad-spectrum vaccine effective against Virus X given: our current knowledge of the virome without any additional discovery, $p_{BSV\,|\,\neg VDS}$; if we discovered 3 times as many viruses as today, $p_{BSV\,|\,3x\,dis}$; and if we discovered the complete virome; $p_{BSV\,|\,full\,virome}$. To calculate the change in likelihood of a BSV given 3x viral discovery, $\Delta p_{BSV|VDS}$; we took the difference between participants answers for $p_{BSV\,|\,full\,virome}$ and $p_{BSV\,|\,\neg VDS.}$

$$\Delta p_{BSV\,|\,VDS} = p_{BSV\,|\,3x\,dis} - p_{BSV\,|\,\neg\,VDS} = 0.084$$

Similarly, to evaluate the change in likelihood of a BSV given discovery of the complete virome, $\Delta p_{BSV|full\,VDS}$; we took the difference between participants answers for $p_{BSV\,|\,full\,virome}$ and $p_{BSV\,|\,\neg VDS.}$

$$\Delta p_{BSV\,|\,full\,VDS} = p_{BSV\,|\,full\,virome} - p_{BSV\,|\,\neg\,VDS} = 0.17$$

##### 3x Viral Discovery

Point estimate using the mean value of each parameter.

$$\begin{aligned} E\,[Harm_{VDS}] &= E\,[Harm_{base}] \times (1 - \Delta p_{BSV\,|\,VDS} \times m_{BSV}) \times (1 - \Delta p_{NPI\,|\,VDS} \times \Delta r_{NPI\,|\,VDS}) \\ &= 7,776,000 \times (1 - 0.09 \times 0.416) \times (1 - 0.14 \times 0.38) \\ &= 7,776,000 \times 0.9113 \\ &= 7,086,672 \end{aligned}$$

$$\begin{aligned} E\,[Benefits_{VDS}] &= E[Harm_{base}] - E[Harm_{VDS}] \\ &= 7,776,000 - 7,086,672 \\ &= 689,328 \end{aligned}$$

**Uncertainty Quantification**

The calculations above use the mean as point estimates for parameters based on survey data, though there is a large amount of uncertainty amongst experts reflected in the wide distributions and confidence intervals of the various questions. To account for uncertainties in our parameter estimates, we conducted Monte Carlo simulations using Python with the NumPy and SciPy libraries. We performed 100,000 iterations, drawing parameter values directly from survey data, from beta distributions derived from our survey data and literature review. For each iteration, we calculated the harm reduction and deaths averted using our model equations, generating distributions of possible outcomes. From these distributions, we computed means, medians, and 90% confidence intervals to characterize the central tendencies and uncertainties in our results. These simulations resulted in a mean harm reduction of approximately 9% and a median of approximately 6.3%, averting approximately 740,911 and 492,307 deaths respectively. Our simulations reveal substantial uncertainty in these estimates, as the 90% central interval for harm reduction spans from 0% to 23.3% translating to a range of 0 to 1.46 million deaths potentially averted over the next decade due to VDS. This wide interval underscores the significant variability in potential outcomes of VDS efforts, and the considerable uncertainty and lack of consensus surrounding the impacts of such interventions.

**Scenario Analysis**

In the initial model, we make the assumption that the change in likelihood of NPIs being targeted towards preventing Virus X, $\Delta p_{NPI\,|\,VDS}$; is equivalent to the change in likelihood of a broad-spectrum vaccine being approved for use against virus X, $\Delta p_{BSV\,|\,VDS}$. Here, we relax that assumption and consider scenarios where they are not equivalent, and run MC simulations setting $\Delta p_{BSV\,|\,VDS}$ to 15%, 30% and 50%, running 100,000 simulations at each level.

Results of MC Simulations at Various Levels of $\Delta p_{NPI\,|\,VDS}$ (likelihood of NPIs being funded through VDS)

| $\Delta p_{NPI\,|\,VDS}$ | Lives Saved |
|---|---|
| 15% | x̄: 694,440 (0 ; 1,506,398)<br>M: 699,840 |
| 30% | x̄: 1,129,123 (69,884 ; 2,235,009)<br>M: 1,075,452 |
| 50% | x̄: 1,695,476 (161,741 ; 3,341752)<br>M: 1,555,200 |

#### Complete Virome

Here we estimate the benefits of VDS if all viruses in animals were discovered and sequenced.

$$E\,[Harm_{full\,VDS}] = E\,[Harm_{base}] \times (1 - \Delta p_{BSV\,|\,full\,VDS} \times m_{BSV}) \times (1 - \Delta p_{NPI\,|full\;VDS} \times \Delta r_{NPI\,|\,full\,VDS})$$
$$= 7,776,000 \times (1 - 0.17 \times 0.416) \times (1 - 0.27 \times 0.52)$$
$$= 7,776,000 \times 0.798$$
$$= 6,211,539$$

$$E\,[Benefits_{full\,VDS}] = E[Harm_{base}] - E[Harm_{full\,VDS}]$$
$$= 7,776,000 - 6,211,539$$
$$= 1,564,461$$

Using the averages as point estimates for survey-based parameters, we estimate discovery of the complete virome could save approximately 989,000 lives in expectation over the next decade.

#### Pandemic Virus Identification

We first establish the probability PVI has successfully characterized Virus X, rather than a different pandemic-capable virus, as:

$$p(PVI) = \frac{n}{v} = \frac{n}{65}$$

where $n$ is the number of viruses characterized and $v$ is the number of pandemic-capable viruses that are equally likely to seed a pandemic event.

$$E\,[Harm_{PVI}] = p(PVI) \times E\,[Harm_{base}] \times (1 - p_{TMCM}\,(\Delta m_{TV} + \Delta m_{TT})) \times (1 - p_{NPI\,|\,PVI} \times \Delta r_{NPI\,|\,PVI})$$

(2)

Consider the case where a single pandemic pathogen has been successfully characterized.

$$E\,[Harm_{PVI\,|\,n=1}] = \frac{1}{65} \times 7,776,000 \times (1 - 0.3544\,(0.5 + 0.094)) \times (1 - 0.48 \times 0.517)$$
$$= 119,631 \times 0.59$$
$$= 70,582$$

Comparing the harm to baseline, we obtain:

$$E\,[Benefits_{PVI\,|\,n\,=\,1}] \;=\; 119,631 \;\times\; (1 \;-\; 0.59) \;=\; 49,048 \text{ lives saved}$$

**Uncertainty Quantification**

We assessed the uncertainty in the estimates for the benefits of identifying a single pandemic-capable virus and identifying all pandemic viruses. For the case of a single pandemic virus, we ran 100,000 MC simulations drawing parameter values directly from survey estimates for $p_{TMCM}$, $p_{NPI\,|\,PVI}$ and $\Delta r_{NPI\,|\,PVI}$ parameters. This resulted in a simulation mean and median of 48,911and 47,725 lives saved, with a 90% CI of 10,500 and 93,587 lives saved.

##### Scaling Up PVI

To estimate the benefits of the entire PVI enterprise, we need to consider how the benefits scale for each additional identified virus. One key observation revealed through our followup survey was In our followup survey, we asked participants about the likelihood of sufficient funding being pooled if multiple pandemic viruses were identified. Using the median estimates of participant answers, we plot the likelihood of acquiring sufficient funds against the number of viruses successfully identified (Fig.4)

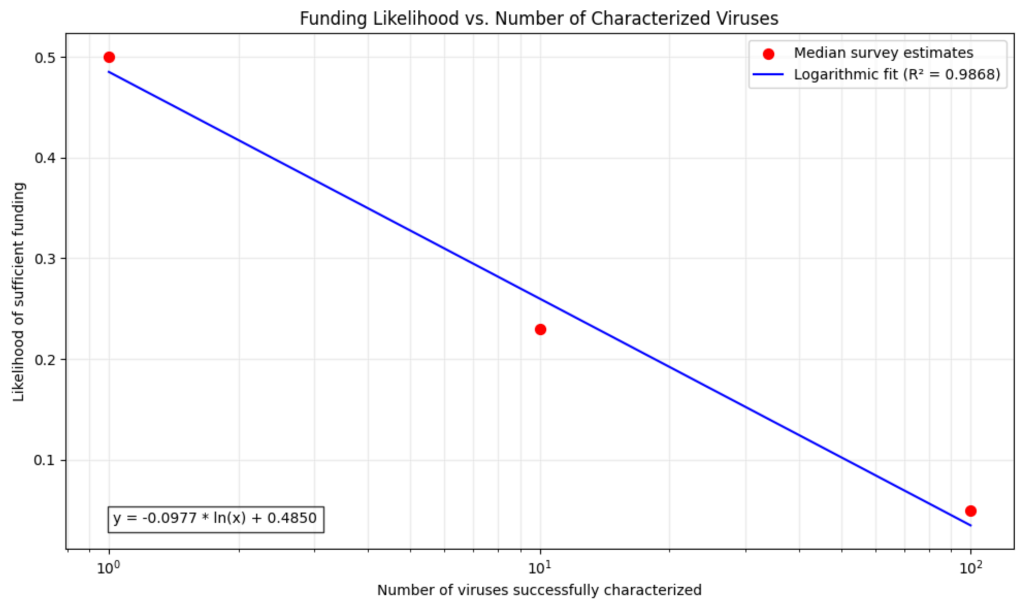

**Supplementary Figure 3 | Relationship between the number of successfully identified pandemic-capable viruses and the likelihood of sufficient funding for countermeasure development.** We fit a logarithmic curve to the data, y = -0.0977ln(x) + 0.485, where y is the likelihood of funding and x is the number of viruses.

In this scaling up version of the model, we assume the probabilities of funding for medical countermeasures and targeted non-pharmaceutical interventions are equal, such that there is a general probability targeted countermeasures will be funded $p_{TMCM}$, where $p_{TCM} = p_{TMCM} = p_{NPI\,|\,PVI}$. We use the logarithmic curve generated by the survey data to generate estimates for $p_{TMCM}$ in the case where multiple pathogens are identified, such that:

$$p_{TCM} = -0.0977\, ln(n) + 0.485$$

We plug this into equation (2), and plot the expected harm in scenarios where PVI and multiple pathogens are identified.

$$E\,[Harm_{PVI\,|\,n}] = \frac{n}{65} \times 7,776,000 \times (1 - p_{TCM}(0.594)) \times (1 - p_{TCM} \times 0.517)$$

Consider the scenario where n = 2

$$E\,[Harm_{PVI\,|\,n\,=\,2}] = \frac{2}{65} \times 7{,}776{,}000 \times (1 - 0.42\,(0.594)) \times (1 - 0.42 \times 0.517)$$
$$= \frac{2}{65} \times 7{,}776{,}000 \times 0.5889$$
$$= 140{,}912$$

$E\,[Benefits_{PVI\,|\,n\,=\,2}] = 239{,}262 - 140{,}912 = 98{,}350$ lives saved

Using this formula, we estimate the number of lives saved based on the number of pandemic-capable viruses identified.

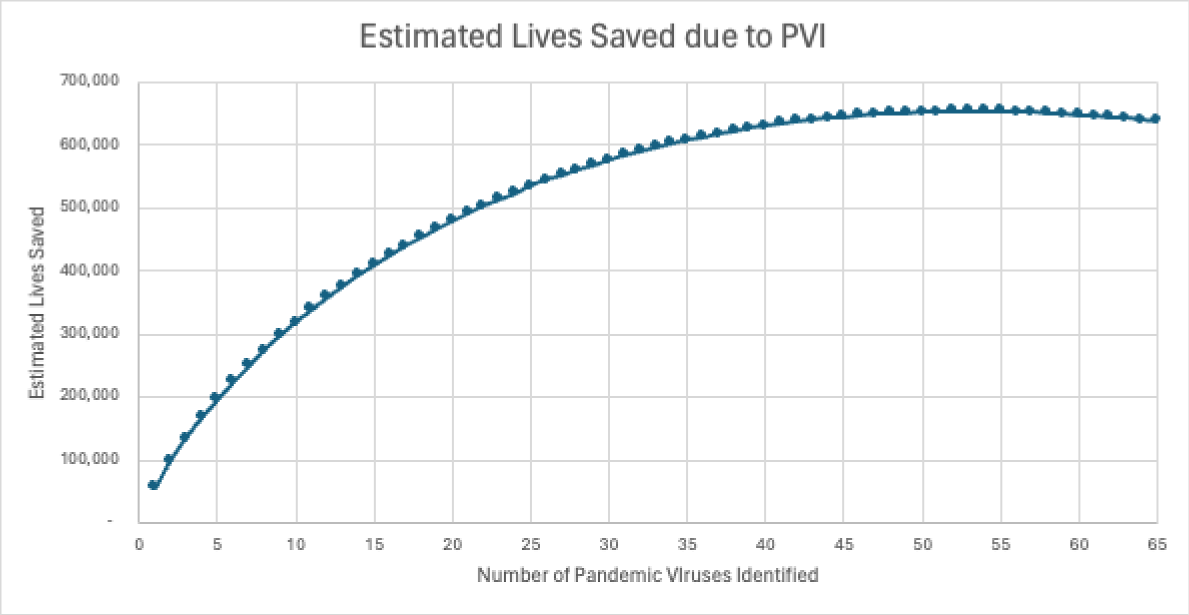

**Supplementary Fig 4 | Benefits of identifying many pandemic-capable viruses.** We estimate if all 65 statistically-equivalent pandemic viruses (an estimated 172 viruses to be characterized in reality) are identified, this would save 642,000 lives in expectation, reducing overall natural pandemic risk by approximately 8% over the next decade.

*Pandemic Virus Identification (PVI)*

Given that pandemic virus identification relies on the characterization of individual pandemic viruses, we first take into consideration the pandemic risk posed by an individual pandemic virus. Through our survey, we estimate there are on average the statistical equivalent of 65 pandemic-capable viruses that are equally likely to seed a pandemic event (172 in reality, with risk concentrated in a minority). It should be noted that to date, no viruses identified as potential pandemic pathogens without spilling over into humans have resulted in the development of targeted interventions. This model starts with the assumption that PVI has successfully characterized a novel zoonotic pandemic-capable virus before spillover, and first estimates the benefits of PVI per successfully identified pandemic virus. We first establish the probability PVI has successfully characterized Virus X, rather than a different pandemic-capable virus, as:

$$p(PVI) = \frac{n}{v} = \frac{n}{65}$$

where $n$ is the number of viruses characterized and $v$ is the number of pandemic-capable viruses that are equally likely to seed a pandemic event.

Based on the key potential benefits noted in literature about PVI, we chose the following parameters to quantify the additional reduction in risk: the likelihood a targeted MCM is funded following identification ( $p_{TMCM}$), the reduction in harm through earlier release of targeted vaccines ($\Delta m_{TV}$) and therapeutics ($\Delta m_{TT}$) due to PVI, the likelihood PVI results in changes to threat-agnostic interventions ($p_{NPI\,|\,PVI}$), and the relative reduction in pandemic risk from PVI informed non-pharmaceutical interventions ($\Delta r_{PVI,\,NPI}$).

We define the reduction in pandemic risk due to PVI as:

$$E\,[Harm_{PVI}] = p(PVI)\ \times E\,[Harm_{base}] \times (1 - p_{TMCM}\,(\Delta m_{TV} + \Delta m_{TT})) \times (1 - p_{NPI\,|\,PVI} \times \Delta r_{NPI\,|\,PVI})$$

(2)

**Incorporating Uncertainty**

The model heavily relies on the likelihood of both medical countermeasures and non-pharmaceutical interventions receiving funding

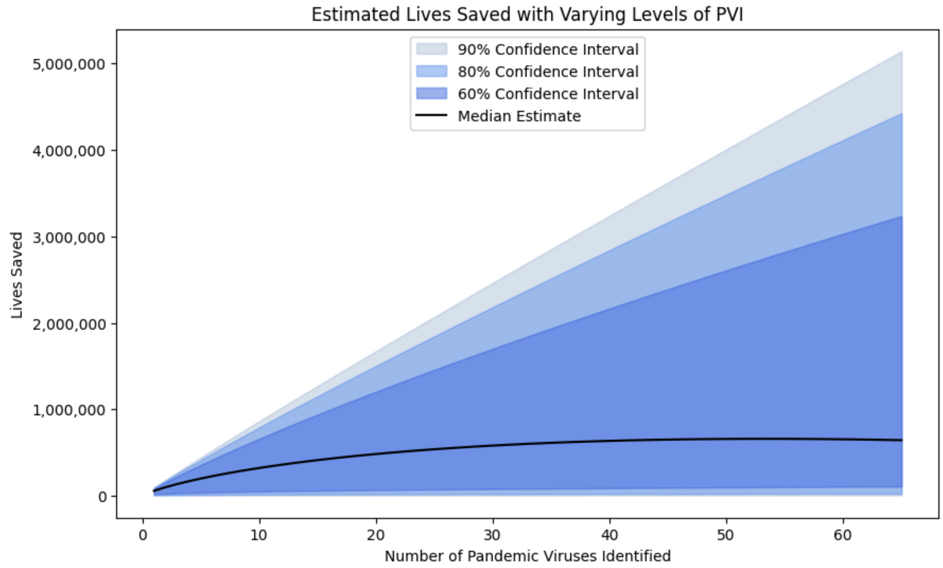

**Supplementary Fig 5 | Estimated number of lives saved as a function of pandemic-capable viruses identified.** The black line represents the model outputs using the median $p_{TMCM}$ (and $p_{NPI\ |\ PVI}$) estimate, and the central interval bands representing model outputs using the 60%, 80% and 90% confidence intervals of the parameter.
